# Bidirectional fibrogenic cross-talk revealed in a human iPSC-derived epithelial-mesenchymal co-culture model of pulmonary fibrosis

**DOI:** 10.64898/2026.01.30.702837

**Authors:** Andrea B Alber, George Kwong, Vishal K Gupta, Porter E Dooley, Jill R Patel, Pushpinder S Bawa, Kasey Minakin, Dakota Jones, Diya Gopal, Huan Souza, Maria Yampolskaya, Eitan Vilker, Chandani Sen, Ansley S Conchola, Pankaj Mehta, Brigitte N Gomperts, Tristan Frum, Jason R Spence, Konstantinos-Dionysios Alysandratos, Darrell N Kotton

## Abstract

Pulmonary fibrosis (PF) can arise from mutations in alveolar epithelial type 2 (AT2) cell-specific genes, but manifests in fibrotic activation of mesenchymal cells, thus involving fibrogenic epithelial-mesenchymal crosstalk. The ligand-receptor interactions underlying the onset and early progression of PF remain poorly understood. Induced pluripotent stem cell (iPSC)-derived models are powerful tools to study respiratory diseases, yet are currently limited to reductionist single lineage epithelial models or multi-lineage systems that lack purity and lung-specificity of the mesenchyme. Here we generate a human iPSC line carrying both a lung mesenchyme-specific reporter (TBX4-LER^tdTomato^) and a reporter for mesenchymal activation/differentiation (ACTA2^GFP^). Applying this line, we develop a directed differentiation protocol capable of generating cells that express key molecular and functional features of primary human developing lung mesenchyme across multiple iPSC genetic backgrounds. We then establish co-cultures of these iPSC-derived lung mesenchymal cells (iLM) with patient-specific iPSC-derived alveolar epithelial type 2 cells (iAT2s) carrying an SFTPC^I73T^ mutation as a model for PF. We find increased expression of fibrotic markers in co-cultures with mutant iAT2s as compared to co-cultures with gene-corrected iAT2s. Moreover, mutant iAT2s express markers of alveolar-basal intermediate (ABI) cells only in the presence of iLM, suggesting that bidirectional crosstalk promotes this aberrant cell state. We identify ligand-receptor pairs enriched in co-cultures with mutant iAT2s, including TGFβ, multiple integrins, and additional genes that have not been previously linked to PF. Finally, we show that small molecule-mediated inhibition of TGFβ or integrins αVβ1/αVβ6 attenuates both fibrotic mesenchymal activation and the presence of ABI cells in iLM/iAT2 co-cultures. Thus, we have established a human iPSC-derived co-culture system that recapitulates key molecular hallmarks of bidirectional fibrogenic epithelial-mesenchymal crosstalk in pulmonary fibrosis, and enables the identification and study of potentially druggable pathways involved in disease initiation and progression.

## Introduction

Pulmonary fibrosis (PF) is a chronic lung disease characterized the presence of aberrant epithelial cells and fibrotic activation of the mesenchyme, resulting in excessive matrix deposition and progressive scarring of the lung^1–5^. Idiopathic pulmonary fibrosis (IPF) remains the most common form of PF, with a median survival of 3-5 years from the time of diagnosis^6–9^. The pathogenesis of PF remains poorly understood, and effective therapies are lacking, as current treatment options mostly target downstream effectors of the disease and only modestly slow down disease progression^10–12^. While animal models and patient tissue samples have allowed the study of later stages of PF^13–16^, research progress has been limited by the lack of human models of disease onset and early progression. PF involves complex cellular crosstalk between multiple lineages, including the epithelium and the mesenchyme^17^. In particular, a broad literature implicates dysfunction of alveolar epithelial type 2 cells (AT2s) in the pathogenesis of PF, including reports of sporadic or familial PF associated with mutations in AT2-specific genes such as *SFTPC*, *SFTPA* and *ABCA3*^18–25^, as well as mutations in telomere-related genes that have been reported to perturb AT2 function based on human and animal models^26–34^. Understanding the mechanisms whereby epithelial-mesenchymal crosstalk leads from epithelial cell-intrinsic dysfunction to the fibrotic activation of the mesenchyme has the potential to open new avenues for therapeutic development.

Complementing animal models and patient tissue samples, induced pluripotent stem cell (iPSC)-derived lineages have emerged as powerful tools to study lung development and respiratory diseases^35,36^. Pluripotent stem cells (PSCs) can differentiate into any lineage of interest through directed differentiation, i.e. the stepwise addition of growth factors and small molecules to recapitulate developmental milestones. Our lab and others have established protocols to generate respiratory epithelial lineages from mouse and human PSCs, including human iPSC-derived alveolar epithelial type 2 cells (iAT2s)^37–49^. iAT2s express key AT2 markers including *SFTPC*, assemble functional lamellar bodies and can be propagated indefinitely without the support of feeder cells^43,50^. We have previously employed patient-derived iAT2s carrying pathogenic variants in the *SFTPC* or *ABCA3* genes to study interstitial lung disease^51–54^. For example, Alysandratos et al. generated patient-derived iAT2s carrying an SFTPC^I73T^ mutation, as well as their gene-edited, corrected counterparts to reveal the epithelial-intrinsic cell dysfunction proposed to occur at the inception of interstitial lung disease^51^. While this model system recapitulated multiple key disease hallmarks, it did not generate alveolar-basal intermediate cells (“ABIs”, also referred to as aberrant basaloid cells, transitional cells, DATP, PATS, ADI, cell cycle arrest, reprogrammed state, or KRT5-/KRT17+ cells), an aberrant epithelial cell state that has been associated with pulmonary fibrosis and in humans expresses a unique gene signature including keratins such as KRT17, but not KRT5^15,16,55–62^. Furthermore, this reductionist single lineage model system was limited by the lack of mesenchymal cells, a key effector cell of the disease, and thus did not allow study of how epithelial cell dysfunction results in fibrogenic mesenchymal activation in pulmonary fibrosis.

iPSC-derived co-cultures/organoids containing both epithelial and mesenchymal cells can be generated either by separate derivation and subsequent co-culture, or by “co-development”, i.e. the differentiation of iPSCs into multiple lineages within the same culture dish^37,63–65^. While co-development can generate multi-lineage organoids with 3-dimensional (3D) organization and close juxtaposition between epithelial and mesenchymal cells, it does not allow for control of the ratio between epithelial and mesenchymal cells or for the removal of any non-lineage specific contaminating cells within the organoid. In contrast, separate derivation and subsequent co-culture allows for purification and characterization of all lineages of interest before co-culture, thus reducing potential confounding factors introduced by the presence of non-characterized cells. Separate derivation and subsequent co-culture has been successfully used to generate human iPSC-derived gastrointestinal organoids^66^ and for co-culture of mouse iPSC-derived lung epithelial progenitors with lung mesenchymal progenitors^67^. Furthermore, two previous studies have established co-cultures of human iPSC-derived respiratory epithelial and human iPSC-derived mesenchymal cells^68,69^. However, it is unclear whether the iPSC-derived mesenchymal cells used in these studies had a lung-specific identity. Importantly, recent single cell RNA sequencing (scRNAseq) studies found specific transcriptional signatures in mesenchymal cells of different organs, both in the developing embryo and in adults^70–72^, highlighting the importance of achieving lung-specificity of both epithelial and mesenchymal cells in co-culture models in order to faithfully study lung development and model respiratory diseases.

Thus, a key hurdle in establishing iPSC-derived lung epithelial-mesenchymal co-cultures for respiratory disease modeling is the differentiation, characterization, and purification of lung-specific mesenchyme from human iPSCs. Our lab has derived functional lung-specific mesenchyme from mouse iPSCs by directed differentiation toward the lateral plate mesoderm and subsequent stimulation of RA and Hh signaling^67^, using a mouse iPSC line carrying an embryonic lung mesenchyme-specific Tbx4 lung enhancer reporter (Tbx4-LER) that has previously been shown to specifically label embryonic lung mesenchyme *in vivo*^73^. In addition, both Kishimoto et al. and Han et al. have published protocols for the directed differentiation of human iPSCs into lung mesenchyme-like cells without a reporter^70,74^, though the efficiency and purity of these differentiation protocols remain incompletely characterized.

Here we engineer human iPSCs carrying a TBX4 lung enhancer reporter (TBX4-LER^tdTomato^) and establish a lung mesenchyme directed differentiation protocol based on stimulation of RA, Hh and Wnt signaling, yielding an average of 69.3 ± 10.4 % of TBX4-LER^tdTomato^+ cells. This protocol, applied to several iPSC lines derived from different human donors, generates iPSC-derived lung mesenchymal cells (iLM) expressing key molecular and functional features of human primary developing lung mesenchyme. We then establish co-cultures of human iLM with patient-derived SFTPC^I73T^ mutant and syngeneic corrected iAT2s as a model for PF and show that co-culture with mutant iAT2s leads to fibrotic activation of iLM, including upregulation of a panel of 5 key fibrotic markers (ACTA2, COL1A1, CTHRC1, FN1 and CTGF). Furthermore, SFTPC^I73T^ mutant iAT2s activate expression of ABI markers only when co-cultured with iLM, suggesting that crosstalk with the mesenchyme is required for the acquisition of the ABI cell state. Finally, we use computational analysis to identify ligand-receptor pairs enriched in mutant iAT2 co-cultures as potential drug targets and demonstrate, using inhibition of TGFβ and integrins αVβ1/αVβ6, how our co-culture system can serve as a platform for drug development.

## Results

### Human iPSC-derived lung mesenchyme (iLM) can be generated *in vitro* by stimulation of RA, Hh and Wnt signaling

To generate a human iPSC reporter line able to track and purify human embryonic lung mesenchyme we engineered a reporter construct consisting of the human TBX4 lung enhancer region^75^ preceding a minimal promoter driving tdTomato (TBX4-LER^tdTomato^, Figure 1a, Supplemental Figure 1a). The TBX4 lung enhancer region is highly conserved between air-breathing species including mouse and human^75^, and activation of the mouse Tbx4 lung enhancer has been shown to be highly specific to the embryonic lung mesenchyme in a triple transgenic Tbx4-LER-rtTA; TetO-Cre; mTmG reporter mouse^73^. In addition, we demonstrated previously that iPSCs generated from this reporter mouse can be used to track the acquisition of the lung mesenchymal progenitor state *in vitro* during directed differentiation^67^. To facilitate tracking of putative human iPSC-derived lung mesenchyme as well as the activation or differentiation state of mesenchymal cells we engineered the TBX4-LER^tdTomato^ reporter construct into human iPSCs, targeting the endogenous AAVS1 locus in human iPSCs, in combination with our previously published endogenous ACTA2^GFP^ reporter^76^, thus generating a bifluorescent reporter iPSC line (BU3 ACTA2^GFP^/TBX4-LER^tdTomato^; hereafter BU3 A2GTT; Figure 1a). After targeting, this line remained karyotypically normal and expressed pluripotency markers (Supplemental Figure 1a-c).

**Figure 1:**
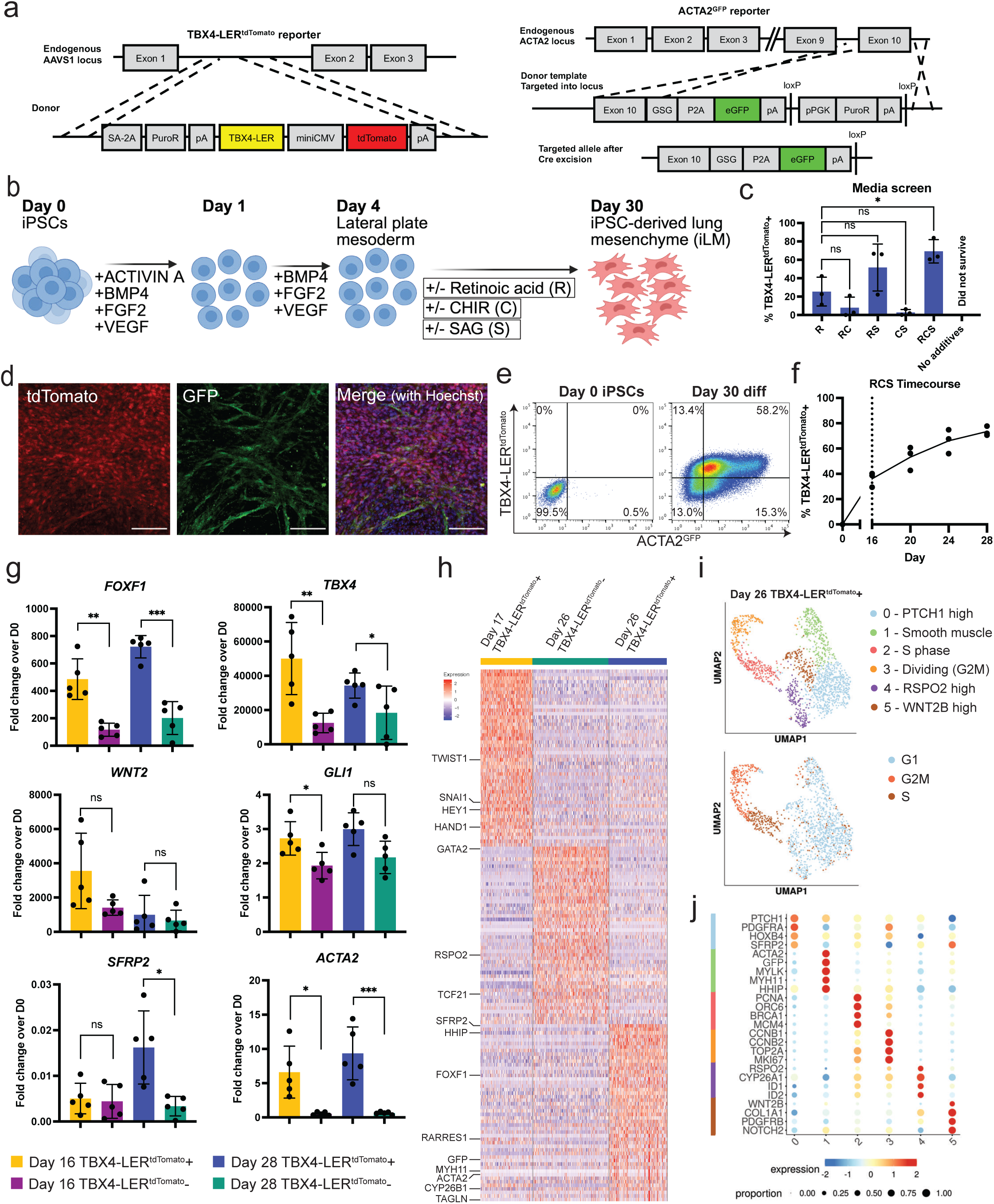
Directed differentiation of human iPSCs into embryonic lung mesenchyme. **a:** Schematic representation of TBX4-LER^tdTomato^ and ACTA2^GFP^ reporter constructs targeted to the indicated endogenous gene loci to generate a bifluorescent mesenchymal reporter iPSC line (BU3 A2GTT). **b**: Lung mesenchyme directed differentiation protocol. To stimulate differentiation into lung mesenchymal progenitors, the combinatorial effects of media supplementation with Retinoic acid (R), CHIR (C), and/or SAG (S) were tested. **c**: Lung mesenchyme directed differentiation media screen, using serum-free medium supplemented with Retinoic acid (R), CHIR (C), and/or SAG (S). TBX4-LER^tdTomato^ reporter fluorescence was determined by flow cytometry on day 30 of differentiation. Each dot represents an independent differentiation. N = 3, bars represent mean and standard deviation. * p<0.05, ns = not significant, as determined by one-way ANOVA. **d**: tdTomato and GFP immunofluorescence microscopy of day 30 cells differentiated in RCS medium. Scale bars = 100 μm. Hoechst = nuclei. **e**: Flow cytometry quantification of day 30 TBX4-LER^tdTomato^/ACTA2^GFP^ reporter cells differentiated in RCS medium, as compared to undifferentiated TBX4-LER^tdTomato^/ACTA2^GFP^ reporter iPSCs. **f**: Temporal dynamics of reporter activation, using RCS medium. tdTomato fluorescence was assessed every 4 days. N = 3 independent differentiations. **g**: RT-qPCR data showing fold change expression relative to day 0 iPSCs of embryonic lung mesenchyme markers of interest in TBX4-LER^tdTomato^ + and – cells on day 16 and 28 of differentiation in RCS medium. N = 5 independent differentiations. Bars represent mean ± SD, * p<0.05, ** p<0.01, *** p<0.001, ns = not significant, as determined by paired, two-tailed t test. **h**: Heatmap showing expression of top 50 differentially expressed genes (DEGs) in day 17 TBX4-LER^tdTomato^ +, day 26 TBX4-LER^tdTomato^ + and day 26 TBX4-LER^tdTomato^ – cells differentiated in RCS medium, as captured by scRNAseq. Genes of interested are highlighted. **i**: Louvain clustering (res 0.5) and cell cycle phase overlay of day 26 TBX4-LER^tdTomato^ + cells, as captured by scRNAseq. **j**: Expression of genes of interest selected from top 50 DEGs in each Louvain cluster.

We generated human iPSC-derived lateral plate mesoderm using our previously published serum-free directed differentiation protocol^76^ and observed expression of the lateral plate mesoderm marker KDR, but no activation of the TBX4-LER^tdTomato^ reporter on day 4 of differentiation (Figure 1b, Supplemental Figure 1d-e). Subsequent culture in serum-free medium containing 2 μM Retinoic acid (RA) yielded an average of 25.5 ± 12.8 % of TBX4-LER^tdTomato^+ cells on day 28 of differentiation (Figure 1b-c), in line with our previous findings identifying RA as a key factor necessary to specify lung mesenchyme from lateral plate mesoderm in our mouse iPSC directed differentiation protocol^67^. Adding both the Wnt agonist CHIR99021 and the Hedgehog agonist SAG to this RA-containing medium (further referred to as “RCS medium”) significantly increased this percentage, yielding an average of 69.3 ± 10.4 % of TBX4-LER^tdTomato^+ human cells by day 28 of differentiation (Figure 1c-e). In contrast, only adding either CHIR99021 or SAG to RA-containing medium (“RC” or “RS” medium) did not significantly alter the TBX4-LER^tdTomato^ percentage, and culture of day 4 cells in serum-free medium without any small molecules led to poor survival and no detectable tdTomato fluorescence (Figure 1c). We profiled the temporal dynamics of TBX4-LER^tdTomato^ reporter activation in RCS medium and found that TBX4-LER^tdTomato^+ cells started to appear by day 16 of differentiation and tdTomato fluorescence gradually increased until day 28 of differentiation (Figure 1f). ACTA2^GFP^ fluorescence also increased over time, leading to a maximum number of TBX4-LER^tdTomato^+/ ACTA2^GFP^+ cells on day 28 (Supplemental Figure 1f).

We profiled the expression of known embryonic lung mesenchyme markers in iPSC-derived TBX4-LER^tdTomato^+ cells by RT-qPCR, immunofluorescence microscopy, and scRNAseq at an early (day 16 for RT-qPCR and day 17 for scRNAseq, respectively) and late (day 28 for RT-qPCR and day 26 for scRNAseq, respectively) stage of differentiation. Day 28 TBX4-LER^tdTomato^+ cells expressed *FOXF1* transcript and FOXF1 protein, a marker known to be expressed throughout the developing embryonic lung mesenchyme beginning at the lateral plate mesodermal stage (Figure 1g, Supplemental Figure 2a). Moreover, *FOXF1* transcript expression was significantly enriched in TBX4-LER^tdTomato^+ compared to TBX4-LER^tdTomato^- cells both on day 16 and 28 of differentiation (Figure 1g). While no individual marker is entirely specific for lung mesenchyme, iPSC-derived cells also expressed a constellation of embryonic lung mesenchyme markers, including *TBX4*, *WNT2*, and *Hh* targets such as *GLI1*, and the expression of multiple markers (*FOXF1*, *TBX4*, *GLI1*, *SFRP2*) was significantly higher in TBX4-LER^tdTomato^+ compared to TBX4-LER^tdTomato^- cells, suggesting that the TBX4 lung enhancer reporter can enrich for lung mesenchyme-like cells (Figure 1g). scRNAseq of day 17 TBX4-LER^tdTomato^+ cells, day 26 TBX4-LER^tdTomato^+ cells and day 26 TBX4-LER^tdTomato^- cells revealed that day 26 TBX4-LER^tdTomato^+ cells were enriched for *FOXF1*, targets of the RA and Hh pathways (*HHIP*, *CYP26B*), and *SFRP2*, which has been described by He et al. as highly expressed in early human lung fibroblasts^77^ (Figure 1h, Supplemental Figure 2b, Supplemental Table 1). Consistent with the observed increase in ACTA2^GFP^ reporter fluorescence, day 26 TBX4-LER^tdTomato^+ cells were also enriched for smooth muscle markers (*ACTA2*, *MYH11*, *TAGLN*), suggesting that this population might include more mature cells. For more detailed understanding of cellular heterogeneity, Louvain clustering analysis revealed that day 26 TBX4-LER^tdTomato^+ cells segregated into 6 distinct clusters, two of which represented G2M and S phase cells, respectively (Figure 1i, j, Supplemental Figure 2c, Supplemental Table 2). The remaining clusters consisted of smooth muscle-like cells (cluster 1), *PTCH1*/*PDGFRA* high cells (cluster 0), *RSPO2* high cells (cluster 4), and *WNT2B* high cells (cluster 5), confirming heterogeneity within the day 26 TBX4-LER^tdTomato^+ population. Of note, similar clusters have been described in primary human fetal lung mesenchyme by Hein et al^78^, including a *PTCH1*/*PDGFRA* high cluster, a smooth muscle cluster, and a *RSPO2* high cluster. While day 17 TBX4-LER^tdTomato^+ cells also expressed multiple key embryonic lung mesenchyme markers, they were enriched for lateral plate mesoderm markers such as *HAND1, KDR, and PRRX1* as compared to 26 TBX4-LER^tdTomato^+ and - cells, suggesting that they might represent an earlier developmental stage (Figure 1h, Supplemental Figure 2d, Supplemental Table 1). Importantly, esophageal/gastric embryonic mesenchymal markers, such as *WNT4*, *MSC* and *BARX1* were expressed at low levels or not expressed in all 3 cell populations (Supplemental Figure 2e). Finally, we tested our differentiation protocol in 3 additional iPSC/embryonic stem cell (ESC) lines with different genetic backgrounds and found similar expression of embryonic lung mesenchymal markers in day 28 cells (Supplemental Figure 2f).

### iLM expresses molecular and functional features of early human fetal lung mesenchyme

To investigate which developmental stage our iPSC-derived mesenchymal cells most closely resemble, we employed two independent methods: i) Single-cell Type Order Parameters (scTOP), a published approach that allows for unbiased and quantitative comparison of PSC-derived cell types to primary cell types using existing scRNAseq atlases as reference datasets, with less susceptibility to batch artifacts than other methods^79,80^; and ii) generation of our own scRNAseq dataset of human fetal lung mesenchyme at 4 different developmental stages between week 8 and 17 post conception. We used scTOP to compare day 17 TBX4-LER^tdTomato^+, day 26 TBX4-LER^tdTomato^+ and day 26 TBX4-LER^tdTomato^- cells to a published scRNAseq atlas of human primary fetal lung mesenchyme from 8 developmental time points between weeks 5 and 22 post conception ^77^ and found that all iPSC-derived samples aligned most strongly with early fibroblasts from week 5 post conception (Figure 2a, Supplemental Figure 3a). In contrast, similarity scores of iPSC-derived cells were lower when compared to other mesenchymal cell types from later stages, including alveolar or adventitial fibroblasts, or non-mesenchymal cell types (Figure 2a, Supplemental Figure 3a). Day 17 TBX4-LER^tdTomato^+, Day 26 TBX4-LER^tdTomato^+ and Day 26 TBX4-LER^tdTomato^- cells were more similar (had higher alignment distributions) to early fibroblasts from week 5 when projected onto multiple cell types, and displayed progressively less overlap with later developmental stages (Figure 2b, Supplemental Figure 3b, c). Furthermore, as predicted from our Louvain clustering analysis (Figure 1i, j), a minor subset of day 26 TBX4-LER^tdTomato^ + cells aligned more closely with “Late Airway Smooth Muscle Cells” (Supplemental Figure 3c). The top 20 most predictive genes for the scTOP alignment scores of day 26 TBX4-LER^tdTomato^+ with early fibroblasts included embryonic lung mesenchyme markers *FOXF1* and *SFRP2*, as well as the RA signaling target gene *CYP26A1*, suggesting that the similarity of our engineered cells to primary early human lung fibroblasts is based on the expression of these key early lung mesenchyme markers (Figure 2c, Supplemental Figure 3d). To compare our iPSC-derived populations to our own scRNAseq dataset of primary human fetal lung mesenchyme from days 57 (week 8), 76 (week 10), 94 (week 13) and 122 (week 17) post conception we performed Louvain clustering of our primary samples and identified a 50-gene set signature differentially expressed at each time point (Supplemental Figure 3e, Supplemental Table 3). Overlaying the 50 gene set onto each of our iPSC-derived samples (Figure 2d), we found that the day 57 differentially expressed gene (DEG) set was expressed the highest of the 4 gene sets, and was enriched in day 26 TBX4-LER^tdTomato^+ cells compared to day 26 TBX4-LER^tdTomato^- cells and day 17 TBX4-LER^tdTomato^+ cells. Taken together, these data suggest that our iPSC-derived cells resemble early primary human fetal lung mesenchyme, with day 26 TBX4-LER^tdTomato^+ cells more closely resembling primary early embryonic cells than TBX4-LER^tdTomato^- cells, displaying heterogeneity and containing a subset of smooth muscle-like cells.

**Figure 2:**
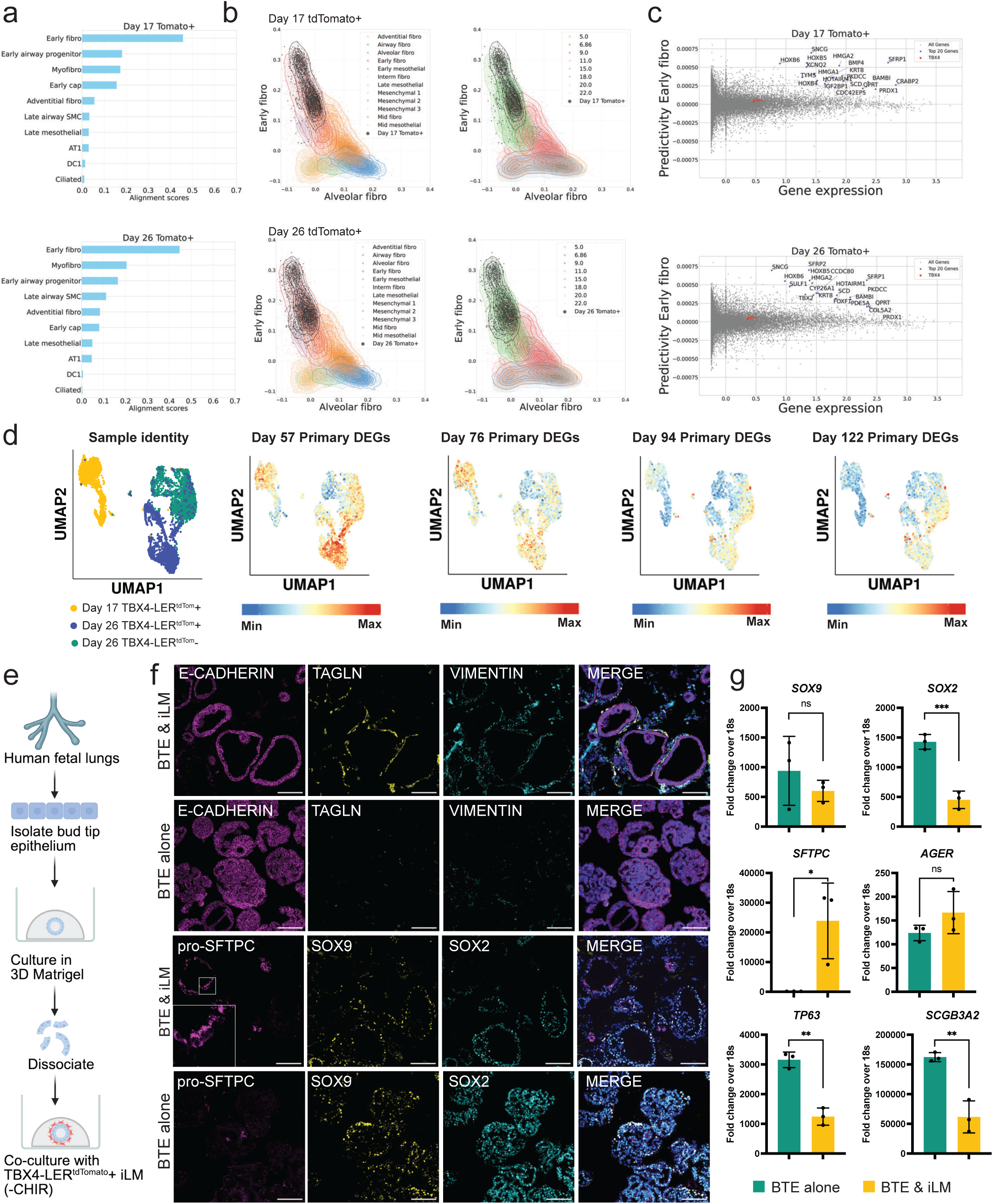
Functional and transcriptomic characterization of human iPSC-derived lung mesenchyme (iLM). **a:** scTOP alignment scores of day 17 TBX4-LER^tdTomato^ + cells and day 26 TBX4-LER^tdTomato^ + cells to selected reference bases of cell types from He et al^77^. **b:** Individual alignment scores for all day 17 TBX4-LER^tdTomato^ + cells and Day 26 TBX4-LER^tdTomato^ + cells and selected primary cells from He et al^77^ plotted against reference bases of Early fibroblasts (y-axis) and Alveolar fibroblasts (x-axis). Primary cells are annotated either by cell type (left panel) or developmental stage in weeks post conception (right panel). **c:** Predictivity scores vs gene expression levels for individual genes, predicting projection of day 17 TBX4-LER^tdTomato^ + cells and day 26 TBX4-LER^tdTomato^ + cells onto the “Early fibroblast” reference basis. The top 20 most predictive genes are highlighted. **d:** UMAP representation of scRNAseq dataset of iPSC-derived day 17 TBX4-LER^tdTomato^ +, day 26 TBX4-LER^tdTomato^ +, and day 26 TBX4-LER^tdTomato^ – cells showing sample identity and expression of gene sets derived from top 50 DEGs in day 57, 75, 94 and 122 primary human fetal lung mesenchyme. **e:** Schematic of experimental approach for co-culture of TBX4-LER^tdTomato^ + iLM with primary human bud tip epithelium (BTE; day 132 post conception). **f:** Immunofluorescence microscopy for E-CADHERIN, TAGLN, VIMENTIN, pro-SFTPC, SOX9 and SOX2 in BTE co-cultured with iLM and BTE cultured alone. Scale bars = 100 μm. **g:** RT-qPCR showing expression of genes of interest in BTE cultured alone and BTE co-cultured with iLM. RT-qPCR was performed on whole wells without re-separation of epithelium and mesenchyme. N = 3 experimental replicates, i.e. separate wells from the same experiment. Bars represent mean ± SD, * p<0.05, ** p<0.01, *** p<0.001, ns = not significant, as determined by unpaired two-tailed student’s t test.

Since iPSC-derived lung mesenchymal cells (hereafter iLM) computationally resemble early human fetal lung mesenchyme, we next evaluated its function by assessing: i) its ability to signal to and interact with developing lung epithelium – a key function of developing lung mesenchyme *in vivo*, and ii) its competence to differentiate to mature mesenchymal lineages. We thus established co-cultures of iLM (day 30 TBX4-LER^tdTomato^+ sorted cells) with primary human embryonic bud tip epithelial progenitors (BTE) from day 132 post conception in 3D Matrigel (Figure 2e). Cells were co-cultured in the absence of CHIR99021, which leads to proximalization of the BTE when BTE is cultured alone^78^. We found that co-cultures self-organized into 3D organoids with closely juxtaposed mesenchyme and epithelium (Figure 2f). Furthermore, co-cultured epithelium expressed higher levels of the distal marker *SFTPC* compared to epithelium cultured alone, as well as significantly lower levels of proximal markers such as *SOX2*, *TP63* and *SCGB1A1* (Figure 2g), suggesting that, similar to primary fetal lung mesenchyme^78^, mesenchymal-epithelial crosstalk from iLM is sufficient to maintain distal identity of BTE.

To test whether iLM is competent to differentiate into more mature mesenchymal lineages, we sorted day 30 TBX4-LER^tdTomato^+ cells and cultured them in RCS medium for an additional 30 days. Day 60 cells significantly decreased expression of the early lung mesenchyme markers *TBX4* and *FOXF1*, and increased expression of smooth muscle markers *ACTA2* and *MYH11* (Supplemental Figure 3f). Furthermore, >95% of cells were ACTA2^GFP^ positive on day 60 (Supplemental Figure 3g), suggesting that iLM is competent to differentiate into the smooth muscle lineage. To determine the competence of iLM cells to differentiate into the endothelial lineage we cultured day 30 TBX4-LER^tdTomato^+ iLM in endothelial differentiation medium. We did not detect expression of the endothelial marker VE-Cadherin after 3 days (Supplemental Figure 3h). In contrast, 26.6 % of day 4 lateral plate mesoderm expressed VE-Cadherin after 3 days of culture in endothelial medium, demonstrating endothelial competence at this earlier developmental stage. These results are in line with previous studies showing that mouse Tbx4-LER^tdTomato^+ lung mesenchymal progenitors lose endothelial competence early in development, both *in vivo* and *in vitro*^67,73^.

### iLM can be co-cultured with iPSC-derived alveolar epithelial type 2 cells (iAT2s) in a 2D submerged co-culture format

We next sought to establish co-cultures of iLM with iAT2s in order to study epithelial-mesenchymal crosstalk in pulmonary disease. To observe meaningful crosstalk, we aimed to achieve survival and outgrowth of both iLM and iAT2s, as well as close juxtaposition between the two lineages. We plated a layer of day 30 TBX4-LER^tdTomato^+ sorted iLM in a well coated with 2D Matrigel and cultured iLM in RCS medium for 2 days. We subsequently plated wildtype iAT2s carrying an SFTPC^tdTomato^ reporter^51^ on top of the mesenchymal cell layer, and co-cultured both lineages for 7 days in iAT2-optimized “CKDCI” medium^43^ (Figure 3a). Co-cultured iLM and iAT2s formed two closely juxtaposed layers (Figure 3b, Supplemental Figure 4a, b). Flow cytometry analysis for EPCAM expression revealed that, after 7 days of co-culture, 70 % of cells were mesenchymal, suggesting survival/outgrowth of mesenchymal cells in these culture conditions (Supplemental Figure 4c). The percentage of tdTomato+ among EPCAM+ cells (i.e. SFTPC^tdTomato^+ epithelial cells), as well as mean fluorescence intensity (MFI) of EPCAM+/tdTomato+ cells was not significantly altered in iAT2s co-cultured in 2D compared to iAT2s cultured alone in 2D (Supplemental Figure 4c). Furthermore, we did not observe any significant differences in expression of AT2 markers as determined by RT-qPCR on EPCAM+ sorted cells, suggesting that 2D co-culture with iLM does not alter iAT2 differentiation/maturation state (Supplemental Figure 4d). Expression of *COL1A1* and *ACTA2* increased in co-cultured iLM compared to either iLM before co-culture or iLM cultured alone in the same 2D conditions (Supplemental Figure 4e), whereas *TBX4*, *FOXF1* and *WNT2* remained unchanged. Thus, iAT2s can be co-cultured with iLM in a 2D submerged format with close juxtaposition of epithelium and mesenchyme, maintenance of AT2 fate, and augmented expression of a subset of mesenchymal markers.

**Figure 3:**
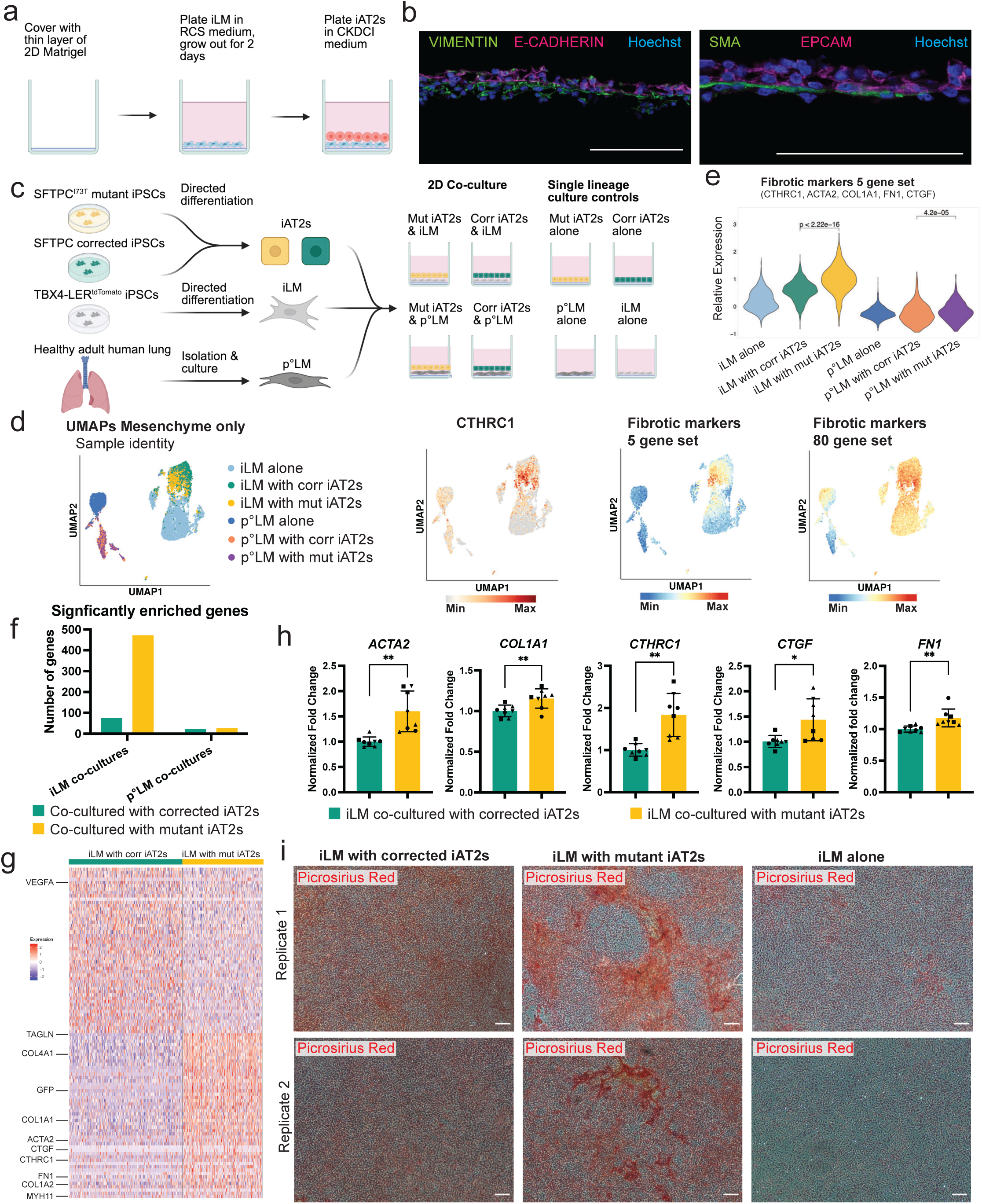
iPSC-derived lung mesenchyme (iLM) co-cultured with patient-derived SFTPC^I73T^ mutant iPSC-derived alveolar epithelial type 2 cells (iAT2s) increases expression of fibrotic markers compared to iLM co-cultured with corrected iAT2s. **a**: Schematic representation of co-cultures of iLM and iAT2s in a 2D submerged co-culture system. **b**: Immunofluorescence staining showing expression of epithelial (E-CADHERIN, EPCAM) and mesenchymal (VIMENTIN, Smooth Muscle Actin [SMA]) markers in cross-sections of 2D submerged iLM/iAT2 co-cultures. Scale bars = 100 μm. Hoechst = nuclei. **c**: Schematic overview of co-cultures captured by scRNAseq. SFTPC^I73T^ mutant (Mut) and corrected (Corr) iPSCs were differentiated into iAT2s and co-cultured with either day 30 TBX4-LER^tdTomato^+ (wild type) iLM or primary human lung mesenchyme isolated and cultured from healthy adult donors. As controls, single lineage 2D cultures of SFTPC^I73T^ mutant or corrected iAT2s, as well as iLM or primary lung mesenchyme were also captured. **d**: scRNAseq of iLM and primary human lung mesenchyme cultured alone or co-cultured with SFTPC^I73T^ mutant or corrected iAT2s. UMAPs represent mesenchyme only, i.e. re-clustered EPCAM- subset of cells, and show sample identity and expression of *CTHRC1*, a gene set of 5 fibrotic markers (*ACTA2*, *COL1A1*, *CTHRC1*, *FN1* and *CTGF*), as well as a larger gene set of 80 markers for fibrotic fibroblasts, generated from Sikkema et al^84^. **e**: Violin plot representation of expression of the 5 fibrotic marker gene set in iLM or primary human lung mesenchyme cultured alone or co-cultured with SFTPC^I73T^ mutant or corrected iAT2s. **f**: Number of significantly enriched genes in iLM co-cultured with SFTPC^I73T^ mutant iAT2s vs iLM co-cultured with corrected iAT2s, as well as in primary lung mesenchyme co-cultured with SFTPC^I73T^ mutant iAT2s vs primary lung mesenchyme co-cultured with corrected iAT2s. **g:** Top 50 DEGs between iLM co-cultured with SFTPC^I73T^ mutant iAT2s and iLM co-cultured with SFTPC^I73T^ corrected iAT2s. Genes of interest are highlighted. **h:** RT-qPCR fold change expression (normalized to iLM co-cultured with corrected iAT2s) of fibrotic markers in re-sorted iLM (EPCAM- cells) after co-culture with SFTPC^I73T^ mutant vs corrected iAT2s. N = 8, symbol shapes indicate experimental replicates, i.e. multiple wells of the same experiment, representative of results from 4 repeated independent experiments performed with iAT2s of varying ages and passages. Bars represent mean ± SD, * p<0.05, ** p<0.01, *** p<0.001, ns = not significant, as determined by paired two-tailed student’s t test. **i:** Picrosirius red stain of iLM co-cultured with SFTPC^I73T^ mutant or corrected iAT2s, or cultured alone for 1 week. Replicates represent two separate experiments. Scale bars = 100 μm.

### Co-cultures of iLM with SFTPC^I73T^ mutant patient-derived iAT2s recapitulate hallmarks of fibrosis

Next, we sought to employ our iLM/iAT2 co-culture system as a disease-relevant in vitro model to study epithelial-mesenchymal crosstalk underlying pulmonary fibrosis. We used an iPSC-line carrying an SFTPC^I73T^ mutation generated from a patient with childhood interstitial lung disease (chILD), a genetic familial form of pulmonary fibrosis, as well as its syngeneic gene-edited/corrected version, both previously generated and published by our lab^51^. We generated iAT2s from both the SFTPC^I73T^ mutant and corrected iPSCs as previously published^43,50^, and established 2D co-cultures with purified wildtype iLM generated from BU3 A2GTT iPSCs. We performed scRNAseq of co-cultures of iLM with SFTPC^I73T^ mutant iAT2s, and iLM with corrected iAT2s, as well as monoculture controls of iLM, SFTPC^I73T^ mutant iAT2s, or corrected iAT2s, each cultured alone in identical conditions (Figure 3c). As a primary cell control, we also included co-cultures of SFTPC^I73T^ mutant and corrected iAT2s with primary human lung fibroblasts isolated from healthy adult donors and expanded in serum-containing culture medium for 7 passages before co-culture. iLM co-cultured with SFTPC^I73T^ mutant iAT2s upregulated expression of known markers of fibrotic activation of fibroblasts^81–83^ (*ACTA2*, *COL1A1*, *CTGF*, *FN1*, *CTHRC1,* both individually and as a 5 gene activation set) as compared to iLM co-cultured with corrected iAT2s (Figure 3d, e, Supplemental Figure 5a, b). We also found increased expression of a gene set of 80 genes found to be enriched in fibrotic fibroblasts, generated from a previously published dataset^84^ (Figure 3d, Supplemental Table 4). In contrast, we observed only weak upregulation of fibrotic markers in primary lung fibroblasts co-cultured with SFTPC^I73T^ mutant iAT2s compared to co-cultures with corrected iAT2s (Figure 3d, e, Supplemental Figure 5a, b). While we found 472 genes to be significantly enriched in iLM co-cultured with SFTPC^I73T^ mutant iAT2s compared to iLM co-cultured with corrected iAT2s, only 26 genes were significantly enriched in primary lung mesenchyme co-cultured with SFTPC^I73T^ mutant iAT2s compared to primary lung mesenchyme co-cultured with corrected iAT2s (Figure 3f, Supplemental Table 5), suggesting that iLM is able to recapitulate fibrogenic mesenchymal activation more reliably than primary lung fibroblasts in this co-culture system. Importantly, many fibrotic markers including *ACTA2*, *COL1A1*, *CTGF*, *FN1*, and *CTHRC1*, several collagens, and smooth muscle markers such as *MYH11* were among the top 50 differentially expressed genes upregulated in iLM co-cultured with SFTPC^I73T^ mutant iAT2s compared to iLM co-cultured with corrected iAT2s (Figure 3g). RT-qPCR on EPCAM- cells after co-culture confirmed that fibrotic markers *ACTA2*, *COL1A1*, *CTGF*, *FN1*, and *CTHRC1* were expressed at significantly higher levels in iLM co-cultured with SFTPC^I73T^ mutant iAT2s compared to iLM co-cultured with corrected iAT2s (Figure 3h). Of note, the percentage of EPCAM- cells after co-culture, but not the percentage of GFP+ mesenchymal cells was significantly increased in co-cultures with SFTPC^I73T^ mutant iAT2s compared to co-cultures with corrected iAT2s (Supplemental Figure 5c). Finally, we used Picrosirius Red staining to assess fibrillar collagen deposition in our co-cultures and detected areas of high staining density in co-cultures with SFTPC^I73T^ mutant iAT2s but not co-cultures with corrected iAT2s or iLM alone (Figure 3i), thus confirming our scRNAseq findings at the protein level.

### SFTPC^I73T^ mutant iAT2s express markers of alveolar-basal intermediate cells (ABIs) only when co-cultured with iLM

We assessed the transcriptomic profiles of SFTPC^I73T^ mutant and corrected iAT2s after co-culture, as well as after iAT2 monoculture under identical 2D culture conditions, by scRNAseq. EPCAM+ cells expressed *NKX2-1* and *SFTPC*, indicating maintenance of AT2 fate after co-culture (Supplemental Figure 5d, Figure 4a). Louvain clustering analysis revealed the emergence of a sub-population of cells (cluster 9) almost exclusively composed of SFTPC^I73T^ mutant iAT2s co-cultured with iLM (Figure 4b-c). This cluster was enriched for *KRT17* and other ABI markers including *KRT7*, *CLDN4*, *SOX4, ANKRD1,* and *LGALS3*, did not express KRT5, and was enriched in a 20-gene module expressed in the KRT5-/KRT17+ epithelial cell population described by Habermann et al. in lungs from patients with PF^15,85^ (Figure 4d, Supplemental Table 6). We identified the top 50 differentially expressed genes between SFTPC^I73T^ mutant iAT2s and corrected iAT2s in iAT2 monocultures and found *SFTPC* to be enriched in SFTPC^I73T^ mutant iAT2s, in line with our previous findings showing higher expression of AT2 maturation genes in SFTPC^I73T^ mutant compared to corrected iAT2s in 3D iAT2 monocultures^51^ (Figure 4e, Supplemental Table 7). Focusing on the co-cultures, we found multiple ABI markers upregulated in the SFTPC^I73T^ mutant iAT2s co-cultured with iLM compared to corrected iAT2s co-cultured with iLM (Figure 4e, Supplemental Table 7). These included *KRT17*, *KRT7*, *GDF15*, *ANKRD1*, *CYR61*, *ITGAV* and *CDKN1A* among the top 50 upregulated genes, and *CLDN4*, *SOX4*, *KRT8*, and *ITGB6* among those significantly upregulated beyond the top 50. Importantly, none of these ABI markers were differentially expressed between SFTPC^I73T^ mutant and corrected iAT2s when cultured alone, suggesting that crosstalk with the iLM is required to drive the emergence of an ABI-like state in this co-culture system. We confirmed by immunofluorescence that KRT17 protein was expressed in co-cultures of SFTPC^I73T^ mutant iAT2s with iLM, but not in co-cultures of corrected iAT2s with iLM or monocultures of either SFTPC^I73T^ mutant or corrected iAT2s (Figure 4f). Of note, KRT17 and other ABI markers were expressed in both SFTPC^I73T^ mutant and corrected iAT2s co-cultured with primary human fetal lung mesenchyme, indicating that co-culture with primary mesenchyme induces an ABI-cell like state independently of the iAT2 genotype (Supplementary Figure 5e-f). This is in agreement with two prior studies reporting loss of the AT2 program when co-culturing iAT2s with primary human lung fibroblast cell lines (MRC5 or AHLM) ^56,85^.

**Figure 4:**
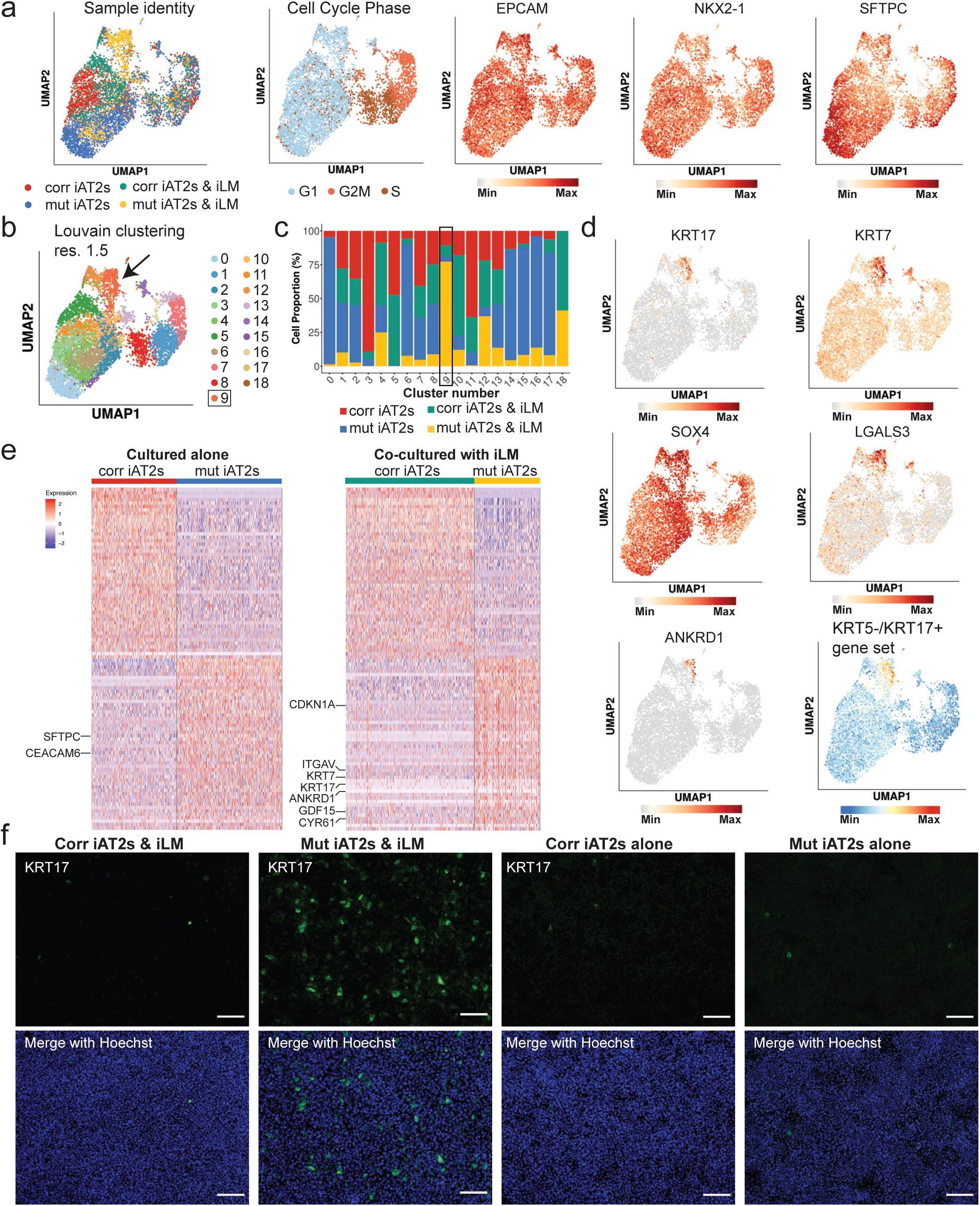
SFTPC^I73T^ mutant iAT2s upregulate markers of alveolar-basal intermediate cells (ABIs) only when co-cultured with iLM. **a:** UMAP representation (EPCAM+ clusters only) of SFTPC^I73T^ mutant vs. corrected iAT2s cultured alone or co-cultured with iLM, showing sample identity, cell cycle phase, and expression of *EPCAM*, *NKX2-1* and *SFTPC*. **b:** Louvain clustering (res 1.5) of EPCAM+ populations. Arrow indicates ABI-like cluster (cluster 9). **c:** Proportion of cells from each sample contributing to each Louvain cluster. ABI-like cluster 9 is highlighted. **d:** UMAP showing expression of ABI markers *KRT17*, *KRT7*, *SOX4*, *LGALS3*, and *ANKRD1*, as well as a gene set derived from KRT5-/KRT17+ cells from Habermann et al^15^, in SFTPC^I73T^ mutant and corrected iAT2s cultured alone or co-cultured with iLM. **e:** Top 50 DEGs between mutant vs corrected iAT2s cultured alone, as well as mutant vs corrected iAT2s co-cultured with iLM. Genes of interest are highlighted. **f:** Immunofluorescence staining showing KRT17 protein expression in SFTPC^I73T^ mutant and corrected iAT2s cultured alone or co-cultured with iLM. Scale bars = 100 μm. Hoechst = nuclei.

### Crosstalk analysis identifies ligand-receptor pairs associated with pulmonary fibrosis

We hypothesized that epithelial-mesenchymal crosstalk could explain both the fibrotic activation of the mesenchyme and the induction of an ABI-like state in co-cultures of iLM with SFTPC^I73T^ mutant iAT2s. To test this, we used NicheNet, a computational method that predicts ligand-receptor interactions based on scRNAseq expression data and prior knowledge of signaling and gene regulatory networks (Figure 5a)^86^. We first compared expression data of iLM cultured alone and iLM co-cultured with SFTPC^I73T^ mutant iAT2s to identify ligands that are specifically upregulated in iLM upon co-culture with SFTPC^I73T^ mutant iAT2s. We found *CTGF* and *TGFB1* to be among the top upregulated ligands (Figure 5b). Both have been described as pro-fibrotic factors expressed by fibroblasts in fibrotic lungs^87,88^, supporting that NicheNet analysis of our iPSC-derived model can identify biologically meaningful factors associated with pulmonary fibrosis. We next performed differential crosstalk analysis to determine the top enriched ligand-receptor pairs in co-cultures with SFTPC^I73T^ mutant vs co-cultures with corrected iAT2s, first defining the iAT2s as sender cells and iLM as receiver cells and then reversing this assignment (Figure 5c, Supplemental Table 8). We identified multiple genes that have been previously described to be associated with pulmonary fibrosis, including *ICAM1*, *LGALS3*, and *AREG* as enriched ligands in SFTPC^I73T^ mutant iAT2 cells^89–91^, as well as *SFRP1* as a mesenchymal ligand^92^, further confirming that our iPSC-derived co-culture system in combination with computational crosstalk analysis can identify ligands suggested by other model systems as mediating fibrotic epithelial-mesenchymal crosstalk. We also identified multiple ligands that have not been previously described in the context of pulmonary fibrosis (Figure 5c, Supplemental Table 8) and thus may provide novel insights into the molecular mechanisms underlying fibrogenic epithelial-mesenchymal crosstalk and potentially serve as druggable targets. Finally, we found multiple integrins, including integrins α1, α5, β1, β6, β5 and αV to be among the top prioritized receptors in SFTPC^I73T^ mutant iAT2s and/or co-cultured iLM, in line with previous reports suggesting a critical role for integrins in PF^93^.

**Figure 5:**
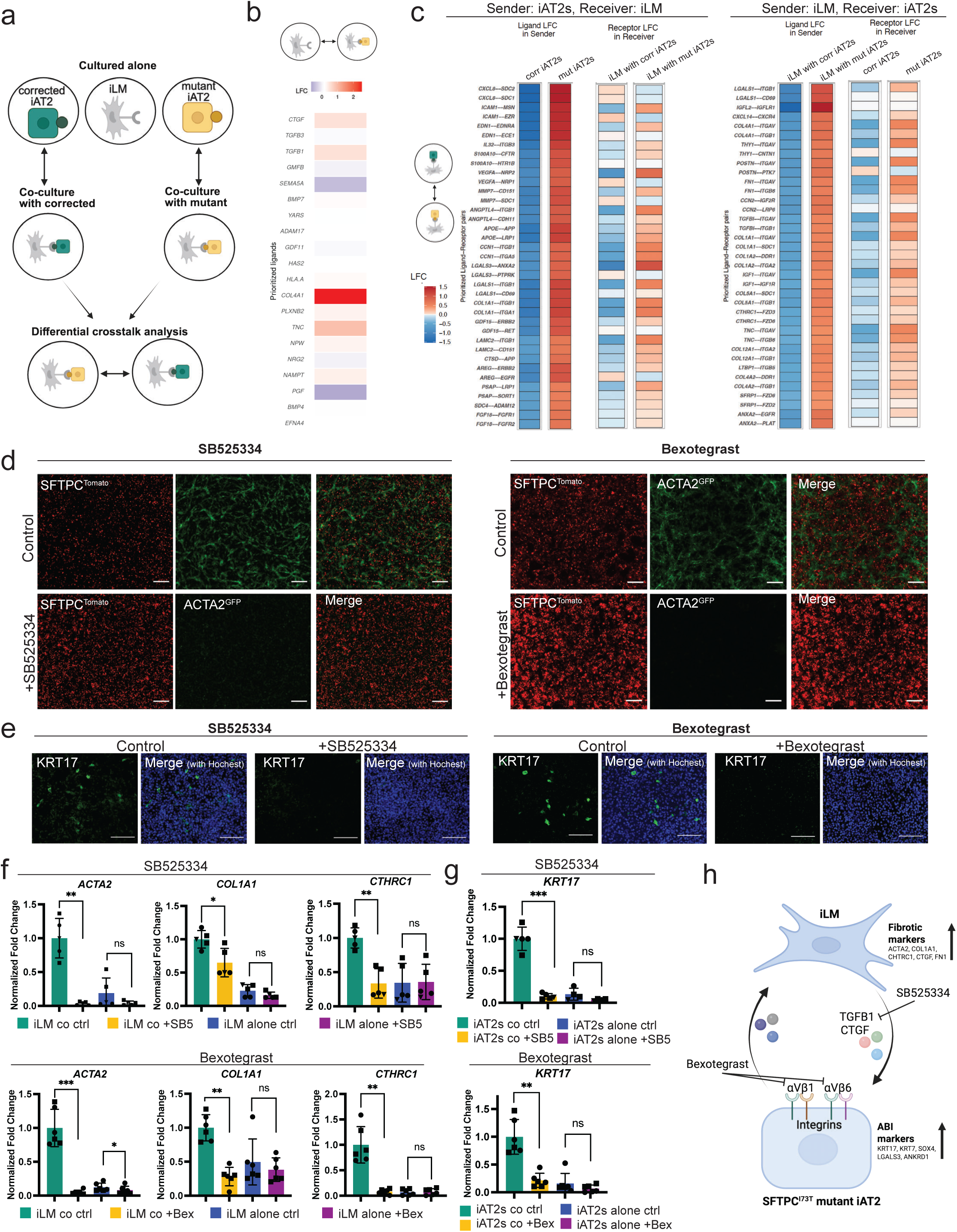
Computational crosstalk analysis identifies ligand-receptor pairs associated with fibrosis. **a:** Schematic representation of computational crosstalk analysis using NicheNet. First, ligands and receptors upregulated upon co-culture (vs single lineage cultures) are identified. Second, ligand-receptor pairs enriched in co-cultures with mutant SFTPC^I73T^ iAT2s (vs corrected iAT2s) are identified. Analysis is performed defining either iLM as sender and iAT2s as receiver, or iAT2s as sender and iLM as receiver. **b:** Expression of top 20 prioritized ligands in iLM co-cultured with SFTPC^I73T^ mutant iAT2s compared to iLM cultured alone. (iLM = sender; SFTPC^I73T^ mutant iAT2s = receiver). **c:** Differential crosstalk analysis. Left panel: Top 20 prioritized ligands in co-cultured SFTPC^I73T^ mutant iAT2s (sender) vs co-cultured corrected iAT2s (sender), with fold change expression of their top prioritized receptors in co-cultured iLM (receiver). Right panel: Top 20 prioritized ligands in iLM co-cultured with SFTPC^I73T^ mutant iAT2s (sender) vs iLM co-cultured with corrected iAT2s (sender), with fold change expression of their top prioritized receptors in SFTPC^I73T^ mutant or corrected co-cultured iAT2s (receiver). LFC = log fold change in gene expression. **d:** SFTPC^tdTomato^ and ACTA2^GFP^ fluorescence in live co-cultures of SFTPC^I73T^ mutant iAT2s and iLM treated with 1 μM SB525334 or 2 μM Bexotegrast for 1 week, as compared to vehicle control (treatment starting on the first day of co-culture). Scale bars = 400 μm. **e:** Immunofluorescence staining for KRT17 protein in co-cultures of SFTPC^I73T^ mutant iAT2s and iLM treated as in (d). Scale bars = 100 μm. Hoechst = nuclei. **f:** RT-qPCR analysis showing fold change expression (normalized to iLM co-cultured control) of fibrotic markers in EPCAM- sorted cells from co-cultures of iLM with SFTPC^I73T^ mutant iAT2s or iLM cultured alone, and treated as in (d). N = 5 for SB525334, N = 6 for Bexotegrast, symbol shapes indicate experimental replicates, i.e. multiple wells of the same experiment. Data points are representative of results from 3 repeated independent experiments for SB525334 and 2 repeated independent experiments for Bexotegrast, performed with iAT2s of varying ages and passages. Bars represent mean ± SD, * p<0.05, ** p<0.01, *** p<0.001, ns = not significant, as determined by paired two-tailed Student’s t test. SB5 = SB525334, Bex = Bexotegrast, ctrl = vehicle control, co = co-cultured, alone = cultured alone. **g:** RT-qPCR analysis showing fold change expression (normalized to iAT2 co-cultured control) of *KRT17* in EPCAM+ sorted cells from SFTPC^I73T^ mutant iAT2s co-cultured with iLM or cultured alone, and treated as in (d). N = 5 for SB525334, N = 6 for Bexotegrast, symbol shapes indicate experimental replicates, i.e. multiple wells of the same experiment. Data points are representative of results from 3 repeated independent experiments for SB525334 and 2 repeated independent experiments for Bexotegrast, performed with iAT2s of varying ages and passages. Bars represent mean ± SD, * p<0.05, ** p<0.01, *** p<0.001, ns = not significant, as determined by paired two-tailed Student’s t test. **h:** Schematic representation of fibrogenic crosstalk between SFTPC^I73T^ mutant iAT2s and iLM.

### iLM/iAT2 co-cultures can be used as platform for drug testing

Having identified candidate pathways underlying fibrogenic epithelial-mesenchymal crosstalk, we next sought to determine whether our iLM/iAT2 co-culture system could serve as a platform for testing drugs, including inhibitors targeting ligands or receptors identified through our NicheNet analysis. We first tested TGFβ inhibition using the ALK5 inhibitor SB525334, which has previously been described to attenuate fibrotic mesenchymal activation in human cells and mice^94,95^. We also tested Bexotegrast, an inhibitor of integrins αVβ1 and αVβ6^96,97^. Given that we identified TGFB1 and CTGF as prioritized ligands in iLM co-cultured with SFTPC^I73T^ mutant iAT2s, integrins αV and β6 as prioritized receptors in SFTPC^I73T^ mutant iAT2s, and integrin β1 as a prioritized receptor in both SFTPC^I73T^ mutant iAT2s and co-cultured iLM (Figure 5b-c), we hypothesized that treatment of co-cultures with SB525334 or Bexotegrast would decrease expression of fibrotic markers in iLM and expression of ABI markers in mutant iAT2s. Treatment of co-cultures of SFTPC^I73T^ mutant iAT2s and iLM for 7 days with either SB525334 or Bexotegrast, compared to vehicle treated controls, resulted in decreased ACTA2^GFP^ fluorescence detectable by live cell microscopy (Figure 5d) and abolished KRT17 protein expression in co-cultured SFTPC^I73T^ mutant iAT2s (Figure 5e). Flow cytometry analysis of ACTA2^GFP^ and SFTPC^tdTomato^ fluorescence in co-cultures versus iLM monocultures confirmed that both SB525334 and Bexotegrast treatment significantly decreased both the percentage and mean fluorescence intensity (MFI) of ACTA2^GFP^+ mesenchymal cells in co-cultured iLM (Supplemental Figure 6a). This effect was less pronounced in SB525334- or Bexotegrast-treated iLM monocultures. In contrast, the SFTPC^tdTomato^ MFI (i.e. the MFI of EPCAM+/tdTomato+ cells) was significantly increased in SB525334 or Bexotegrast-treated co-cultured SFTPC^I73T^ mutant iAT2s (Supplemental Figure 6b). Finally, RT-qPCR analysis after SB525334 or Bexotegrast treatment revealed a significant decrease in the expression of the fibrotic markers *ACTA2*, *COL1A1*, *CTHRC1*, *CTGF* and *FN1* in co-cultured iLM (Figure 5f, Supplemental Figure 6c), consistent with the ACTA2^GFP^ reporter results. Furthermore, expression of the ABI markers *KRT17* and *KRT7* was significantly reduced with treatment in co-cultured SFTPC^I73T^ mutant iAT2s, but not mutant iAT2s cultured alone (Figure 5g, Supplemental Figure 6d). These findings suggest that inhibition of TGFβ and of integrins αVβ1 and αVβ6 indeed decreases both fibrotic activation of the mesenchyme and induction of the ABI-like state in iAT2s (Figure 5h), and that it acts on crosstalk between SFTPC^I73T^ mutant iAT2s and iLM, rather than on either lineage alone. Our findings also demonstrate that our iLM/iAT2 co-culture system can serve as a platform for drug testing in the future, with ACTA2^GFP^ fluorescence and KRT17 staining providing visual readouts suitable for larger scale screening.

## Discussion

In this study we employ directed differentiation of human iPSCs to generate cells expressing key molecular and functional features of early embryonic human lung mesenchyme, including expression of embryonic lung mesenchyme markers, competence to differentiate toward more mature cells, and the ability to signal to and interact with embryonic epithelial lung bud tip progenitors. We engineer iPSCs carrying a lung mesenchyme-specific TBX4 lung enhancer reporter (TBX4-LER), allowing the tracking and quantification of the acquisition of the embryonic lung mesenchymal fate and the successful purification of differentiated cells. We find that simultaneous stimulation of the RA, Hh and Wnt signaling pathways yields the highest percentage of TBX4-LER^tdTomato^+ cells, in line with our previously published protocol for the directed differentiation of mouse iPSCs into lung mesenchyme^67^, and consistent with two previous studies describing approaches for directed differentiation of human iPSCs into lung mesenchyme^70,74^. While transcriptomic analysis using scTOP indicates that both TBX4-LER^tdTomato^+ and TBX4-LER^tdTomato^- cells have similarity with early embryonic lung mesenchymal cells, we show by RT-qPCR (Figure 1g) and scRNAseq (Figure 1h, 2d) that TBX4-LER^tdTomato^+ cells express multiple lung mesenchymal markers at significantly higher levels than TBX4-LER^tdTomato^- cells, suggesting that the TBX4 lung enhancer reporter can enrich for early lung mesenchyme-like cells. Most of our current knowledge on the pathways involved in lung mesenchyme specification *in vivo* stems from animal models, as lung mesenchyme specification in humans has been challenging to study and has mostly relied on temporal snapshots from rare human embryonic tissue samples. Thus, our study provides important insights into the molecular mechanisms of human lung mesenchyme specification.

A key goal of this study was to advance PF modeling beyond existing systems, which are limited to either single epithelial lineages or iPSC-derived organoids generated by co-development that contain multiple lineages but lack purity and controlled epithelial to mesenchymal ratios, limiting their ability to faithfully recapitulate epithelial-mesenchymal crosstalk. We thus establish a 2D human co-culture model of iLM and iAT2s using separate derivation and subsequent co-culture, an approach that enables tight control of the identity and quality of each cell type, as well as the ratio of epithelial to mesenchymal cells used for co-culture. We find close juxtaposition of epithelium and mesenchyme in these co-cultures (Figure 3b), an important feature when aiming to study cellular crosstalk. We demonstrate that co-cultures of iLM with patient-derived iAT2s carrying an SFTPC^I73T^ mutation recapitulate key hallmarks of pulmonary fibrosis, including fibrotic activation of the mesenchyme and expression of ABI markers in the epithelium. Importantly, upregulation of ABI markers is only observed in co-cultured SFTPC^I73T^ mutant iAT2s, whereas these markers are not upregulated in co-cultures with syngeneic corrected iAT2s, or in SFTPC^I73T^ mutant iAT2s cultured alone in the same 2D conditions (Figure 4d-f), or embedded in 3D Matrigel^51^, suggesting that bidirectional crosstalk with the mesenchyme is required for the acquisition of an ABI-like state and highlighting the importance of multilineage model systems for *in vitro* disease modeling. Furthermore, the finding that co-culture of SFTPC^I73T^ mutant iAT2s with primary adult wildtype lung fibroblasts revealed little evidence for fibrotic upregulation compared to co-cultures with iLM (Figure 3d, e), suggests that iLM more reliably recapitulates key disease hallmarks in this co-culture system.

We have used scRNAseq and computational crosstalk analysis by NicheNet^86^ to determine ligands and receptors specifically enriched in co-cultures with SFTPC^I73T^ mutant iAT2s (Figure 5a-c). Of note, our analysis has identified multiple genes that have been previously associated with pulmonary fibrosis, confirming that this approach can reliably identify disease-relevant genes and pathways. Our approach also revealed multiple ligands and receptors that have not yet been linked to pulmonary fibrosis. Future work can focus on testing these candidate genes and investigating whether they might serve as drug targets in the future. We have demonstrated that TGFβ inhibition by SB525334, as well as inhibition of integrins αVβ1 and αVβ6 by Bexotegrast decreases fibrotic activation of the mesenchyme, as well as expression of ABI markers in the epithelium, suggesting that our *in vitro* model system can serve as a platform for drug testing, using fluorescence microscopy readouts based on ACTA2^GFP^ and KRT17 expression that also allow for future larger scale screens. Furthermore, our approach of separate derivation and subsequent co-culture will allow for cell type-specific modulation of genes of interest by CRISPRi or knock-out, which should help to understand the molecular mechanisms underlying fibrogenic epithelial-mesenchymal crosstalk.

In this study we have used patient-derived iPSCs carrying an SFTPC^I73T^ mutation as a model to study epithelial-mesenchymal crosstalk in pulmonary fibrosis. Our iPSC-derived co-culture system will also allow for studying other disease-relevant mutations, including mutations in the Brichos domain of *SFTPC*, mutations in the *ABCA3* and *SFTPB* genes, as well as telomerase-asssociated mutations. This will broaden our understanding of the differences or similarities in ligand/receptor pairs associated with different types of disease-causing mutations and should help to develop appropriate therapies.

### Limitations of the study

While our iPSC-derived lung mesenchyme (iLM) possesses several key molecular and functional features of primary human fetal lung mesenchyme, we also observed some important differences in expression of key lung mesenchymal markers, such as relatively low expression levels of *WNT2*. Furthermore, our iLM remains embryonic-like, even after co-culture with iAT2s. Future work will address how iLM can be matured *in vitro* and instructed to differentiate into mature mesenchymal cell types of interest, such as alveolar or adventitial fibroblasts.

We used co-culture of SFTPC^I73T^ mutant iAT2s with primary adult wildtype lung fibroblasts as a primary control and found fewer differentially expressed genes and less fibrotic activation before vs. after co-culture in primary cells, as compared to co-cultures with iLM (Figure 3d, e), limiting the ability to use primary cells to study fibrogenic cross talk, at least in this model system. Importantly, this need not detract from efforts to develop new alternative model systems that rely on patient-specific or primary lung mesenchyme. For example, the primary human lung mesenchyme used in our study, as in many prior reports, was composed of lung fibroblasts that were cultured and serially passaged in serum-containing medium before co-culture, which likely affects primary cell molecular and fibrogenic phenotype. Lower passage or alternatively purified primary lung fibroblasts, or different co-culture model systems might be less affected and more able to recapitulate fibrotic mesenchymal activation using primary cells. We anticipate our iPSC-derived model system to serve as an important complement to emerging primary cell model systems.

Finally, while our iPSC-derived epithelial-mesenchymal co-culture system contains two key lineages involved in pulmonary fibrosis, it is currently lacking other lineages with suggested roles in the disease, most importantly the immune or endothelial lineages. Further work will be needed to add those lineages to the co-cultures to investigate multi-lineage crosstalk. Despite these limitations, we anticipate the ability to derive human lung-specific mesenchyme either in isolation or for use in co-cultures with lung epithelial lineages, as presented here, provides a novel source of cells representing an additional lung compartment that can be difficult to access in patients and significantly augments the complexity of existing human models of lung development and disease.

---

**Supplemental Figure 1:**
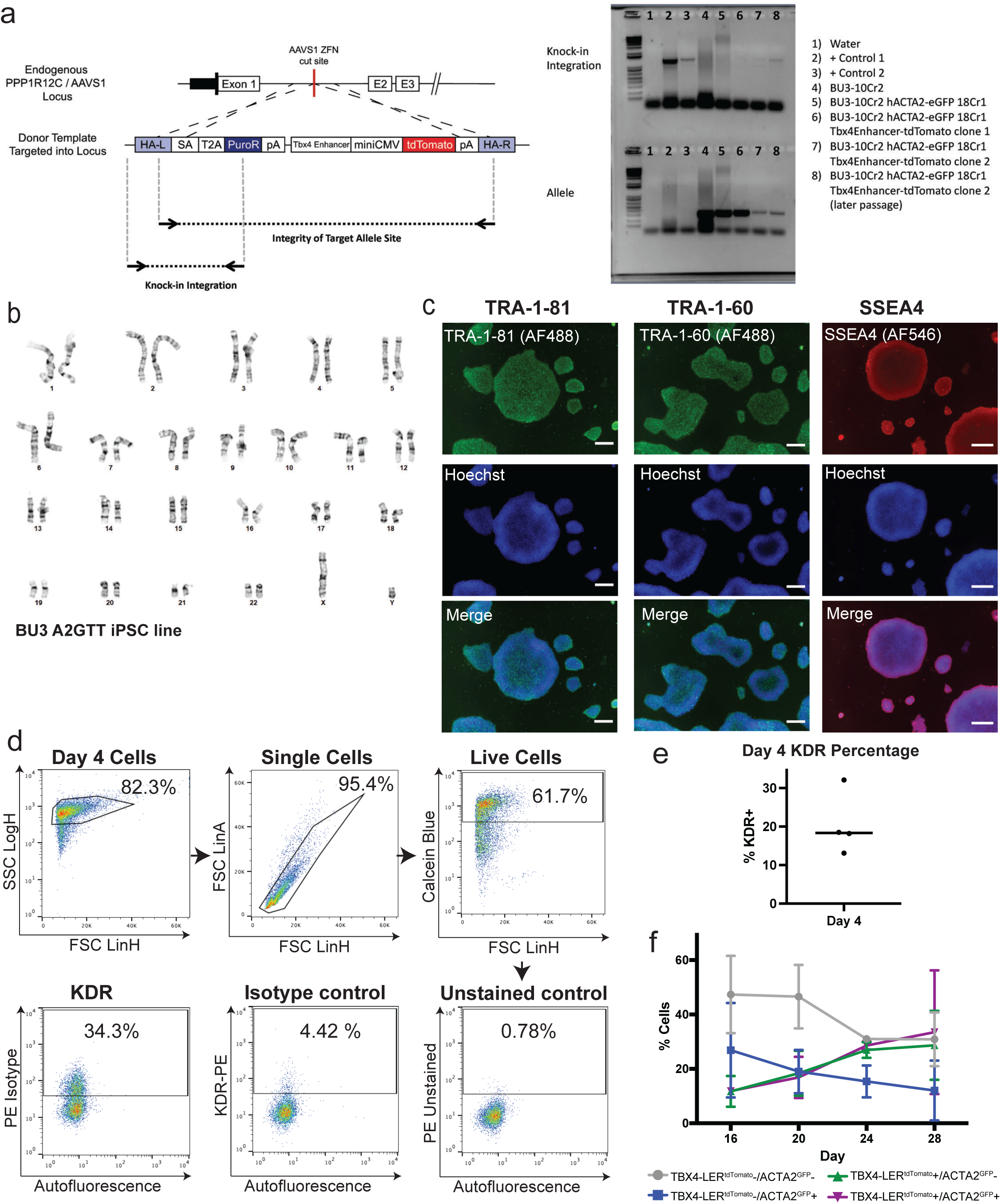
Generation and characterization of a human ACTA2^GFP^/TBX4-LER^tdTomato^ (A2GTT) reporter iPSC line. **a:** Schematic of editing strategy to knock-in a reporter construct consisting of the human TBX4 lung enhancer region preceding a minimal promoter driving tdTomato into the AAVS1 locus. Zinc finger nucleases (ZFNs) were used to target the construct to the endogenous AAVS1 (safe harbor/open) gene locus in iPSCs. Schematic shows two primer sets used to screen iPSC clones for targeted insertion of the TBX4-LER^tdTomato^ reporter construct to the AAVS1 locus: 1) “Knock-in Integration” primer set to show targeting of the reporter construct to the AAVS1 locus; and 2) “Allele Integrity” primer set to determine mono-allelic or bi-allelic integration of the reporter construct. Gel shows results of PCR screening of parental BU3 ACTA2^GFP^ reporter iPSC line (Lane #5) and two clones of ACTA2^GFP^/TBX4-LER^tdTomato^ (BU3 A2GTT) reporter iPSC line ((Lane #6 and Lane#7/8, respectively) with the 2nd clone showing knock-in of the TBX4-LER^tdTomato^ reporter construct with mono-allelic knock-in. **b:** Karyotyping results showing normal karyotype of BU3 A2GTT reporter iPSC line. **c:** Expression of pluripotency markers in BU3 A2GTT reporter iPSC line. Scale bars = 400 μm. Hoechst = nuclei. **d:** Example of gating strategy and representative flow cytometry plots showing KDR stain, isotype and unstained control on day 4 of differentiation (lateral plate mesoderm). **e:** Percentage of KDR+ cells on day 4 of differentiation. N = 4 independent differentiations. **f:** Percentage of TBX4-LER^tdTomato^ and ACTA2^GFP^ reporter positive cells over time. N = 3 independent differentiations, reporter fluorescence quantified by flow cytometry.

**Supplemental Figure 2:**
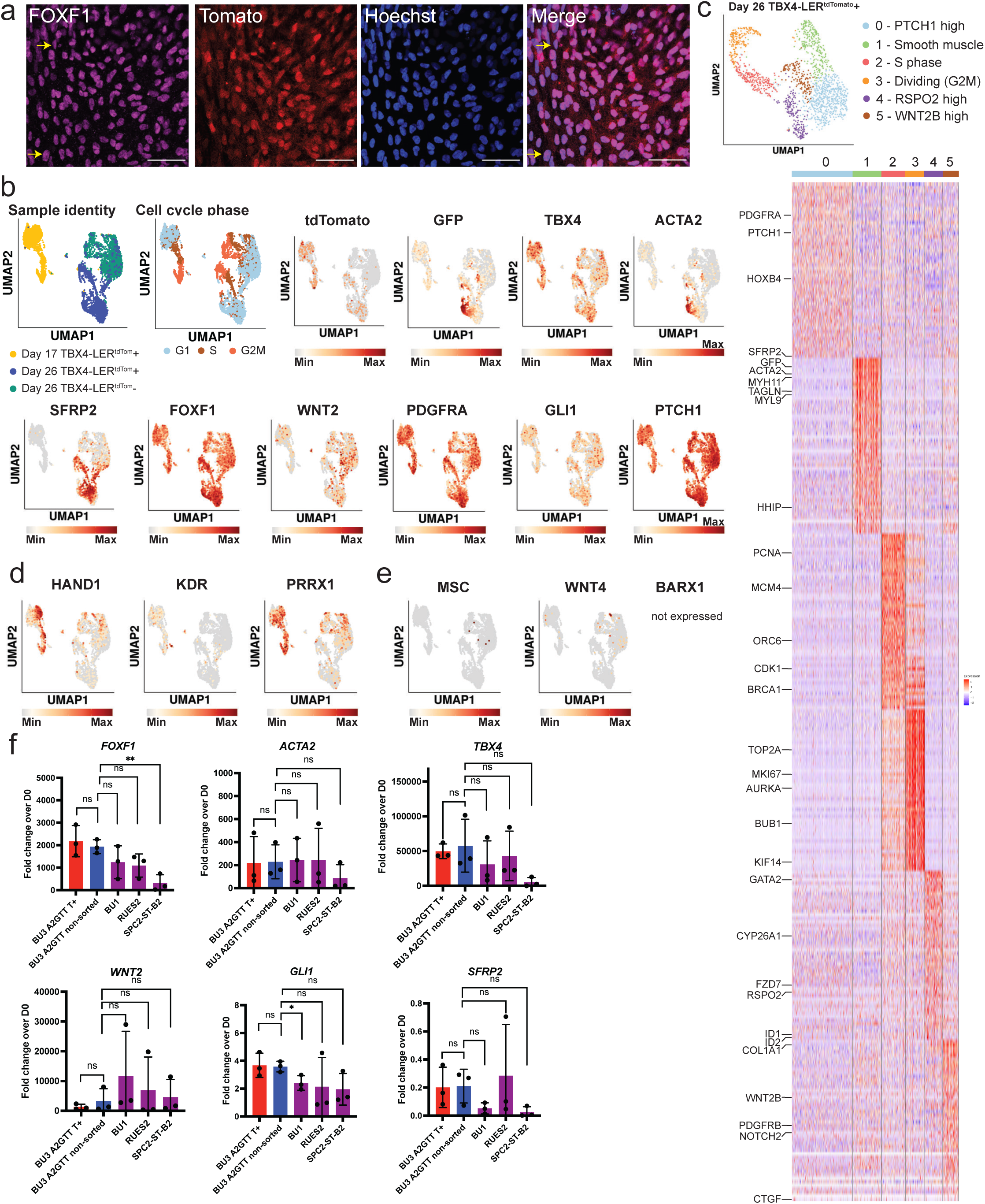
Expression of lung mesenchyme markers in iPSC-derived lung mesenchyme (iLM). **a:** Immunofluorescence showing expression of FOXF1 protein in day 28 BU3 A2GTT reporter cells differentiated in RCS medium (non-sorted). All TBX4-LER^tdTomato^+ cells express FOXF1, whereas a few rare FOXF1+ cells do not express tdTomato (see arrows). Scale bars = 50 μm. Hoechst = nuclei. **b:** UMAP representation of scRNAseq dataset showing sample identity, cell cycle phase overlay and expression of tdTomato, GFP, and lung mesenchyme markers of interest in day 17 TBX4-LER^tdTomato^ +, day 26 TBX4-LER^tdTomato^ + and day 26 TBX4-LER^tdTomato^ – cells. **c**: Top 50 DEGs in Louvain clusters (res 0.5) of day 26 TBX4-LER^tdTomato^ + cells. Genes of interest are highlighted. **d:** UMAP representation showing expression of lateral plate mesoderm markers in day 17 TBX4-LER^tdTomato^ +, day 26 TBX4-LER^tdTomato^ + and day 26 TBX4-LER^Tomato^ – cells. **e:** UMAP representation showing lack of expression of non-lung (gastric/esophageal) embryonic mesenchymal markers in day 17 TBX4-LER^tdTomato^ +, day 26 TBX4-LER^tdTomato^ + and day 26 TBX4-LER^tdTomato^ – cells. **f:** RT-qPCR showing fold change expression relative to day 0 iPSCs of embryonic lung mesenchyme markers of interest in day 28 ACTA2^GFP^/TBX4-LER^tdTomato^ cells sorted for TBX4-LER^tdTomato^ (BU3 A2GTT T+) or non-sorted (BU3 A2GTT non-sorted), as well as non-sorted day 28 iLM derived from 3 other iPSC/ESC lines with different genetic backgrounds. N = 3 independent differentiations. Bars represent mean ± SD, * p<0.05, ** p<0.01, *** p<0.001, ns = not significant, as determined by unpaired two-tailed student’s t test.

**Supplemental Figure 3:**
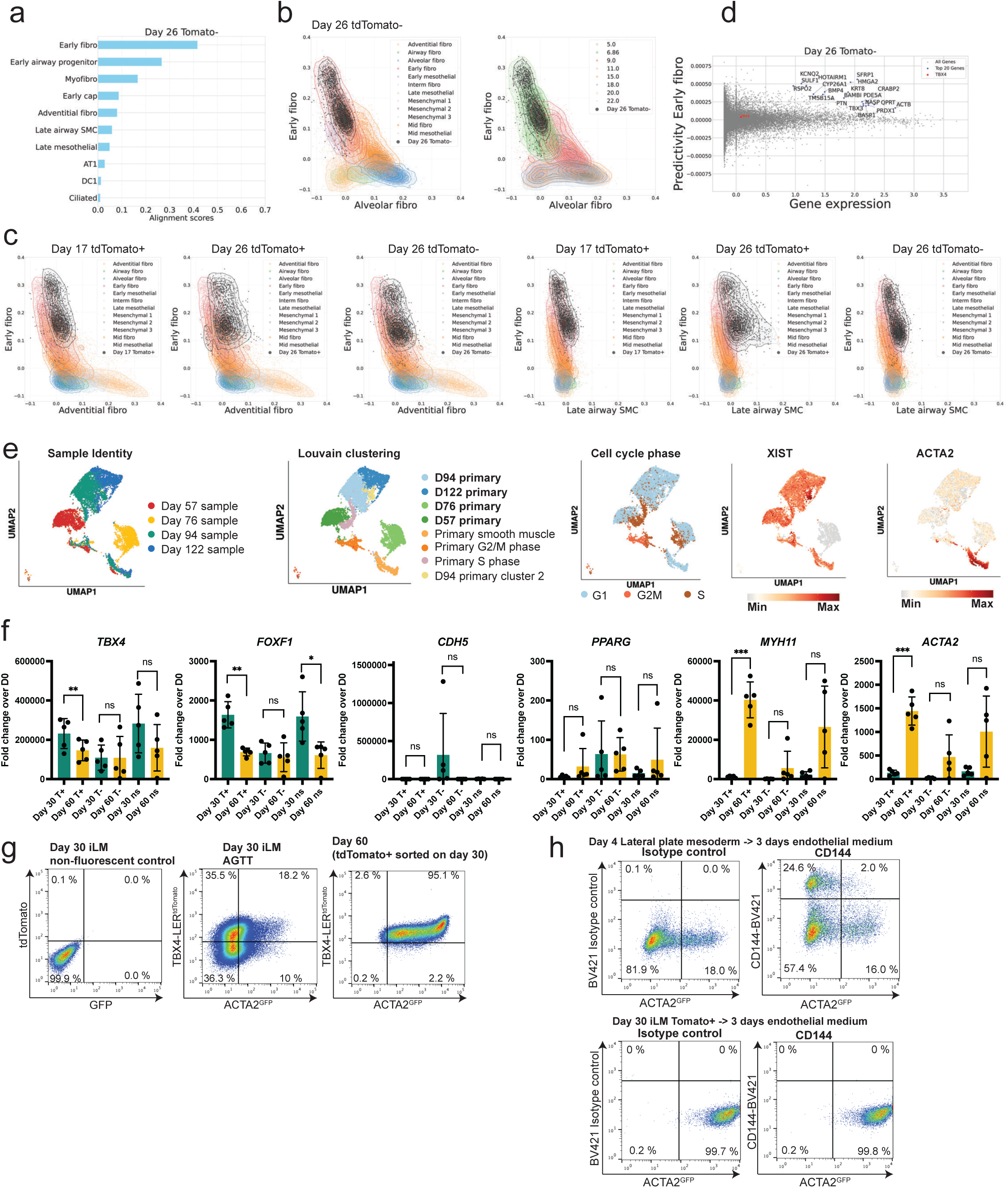
Transcriptomic characterization and developmental competence of human iLM. **a:** scTOP alignment scores of day 26 TBX4-LER^tdTomato^ - cells to selected reference base cell types from He et al^77^. **b:** Individual alignment scores for all day 26 TBX4-LER^tdTomato^ - cells and selected primary cells from reference dataset against early fibroblasts and alveolar fibroblasts. Primary cells are annotated by either cell type (left panel) or developmental stage in weeks post conception (right panel). **c:** Individual alignment scores for all day 17 TBX4-LER^tdTomato^ + cells, day 26 TBX4-LER^tdTomato^ + and 26 TBX4-LER^tdTomato^ - cells and selected primary cells from He et al^77^ plotted against reference bases of early fibroblasts (y-axis) and adventitial fibroblasts (x-axis), or early fibroblasts (y-axis) and late airway smooth muscle cells (x-axis), respectively. Primary cells are annotated by cell type. **d:** Predictivity scores vs gene expression levels for individual genes, predicting projection of day 26 TBX4-LER^tdTomato^ - cells onto the “Early fibroblasts” reference basis. The top 20 most predictive genes are highlighted. **e:** ScRNAseq of primary human fetal lung mesenchyme from days 57, 76, 94 and 122 post conception. UMAPs show sample identity, Louvain clustering (res 0.5), cell cycle phase overlay and expression of *XIST* and *ACTA2*. Top 50 DEGs from the D57 primary, D76 primary, D94 primary and D122 primary Louvain clusters were used to generate gene sets shown in Figure 2d. **f:** RT-qPCR data showing fold change expression over day 0 iPSCs of lung mesenchyme markers and more mature lineage markers in day 30 TBX4-LER^tdTomato^+, TBX4-LER^tdTomato^- and non-sorted (ns) iLM cells, as well as cells that were sorted for TBX4-LER^tdTomato^ +/- or collected on day 30 without sorting, re-plated in RCS medium for an additional 30 days (day 60) and subsequently collected without further sorting. Bars represent mean and standard deviation, N = 5 independent differentiations. Bars represent mean ± SD, * p<0.05, ** p<0.01, *** p<0.001, ns = not significant, as determined by paired two-tailed student’s t test. **g:** Representative flow cytometry plot showing tdTomato and GFP fluorescence in day 30 iLM derived from the TBX4-LER^tdTomato^/ACTA2^GFP^ reporter line and a non-fluorescence control line, as well as day 60 TBX4-LER^tdTomato^/ACTA2^GFP^ reporter cells that were sorted for TBX4-LER^tdTomato^+ on day 30 and re-plated in RCS medium for an additional 30 days. **h:** Endothelial lineage competence of control iPSC-derived lateral plate mesoderm (day 4 cells) vs loss of endothelial competence in day 30 iLM. Endothelial marker CD144 (VE-CADHERIN) cell surface protein expression in iPSCs that were differentiated into lateral plate mesoderm (day 4) and subsequently cultured in endothelial medium for 3 days, as well as iPSCs that were differentiated towards into iLM (day 30), TBX4-LER^tdTomato^+ sorted and subsequently cultured in endothelial medium for 3 days.

**Supplemental Figure 4:**
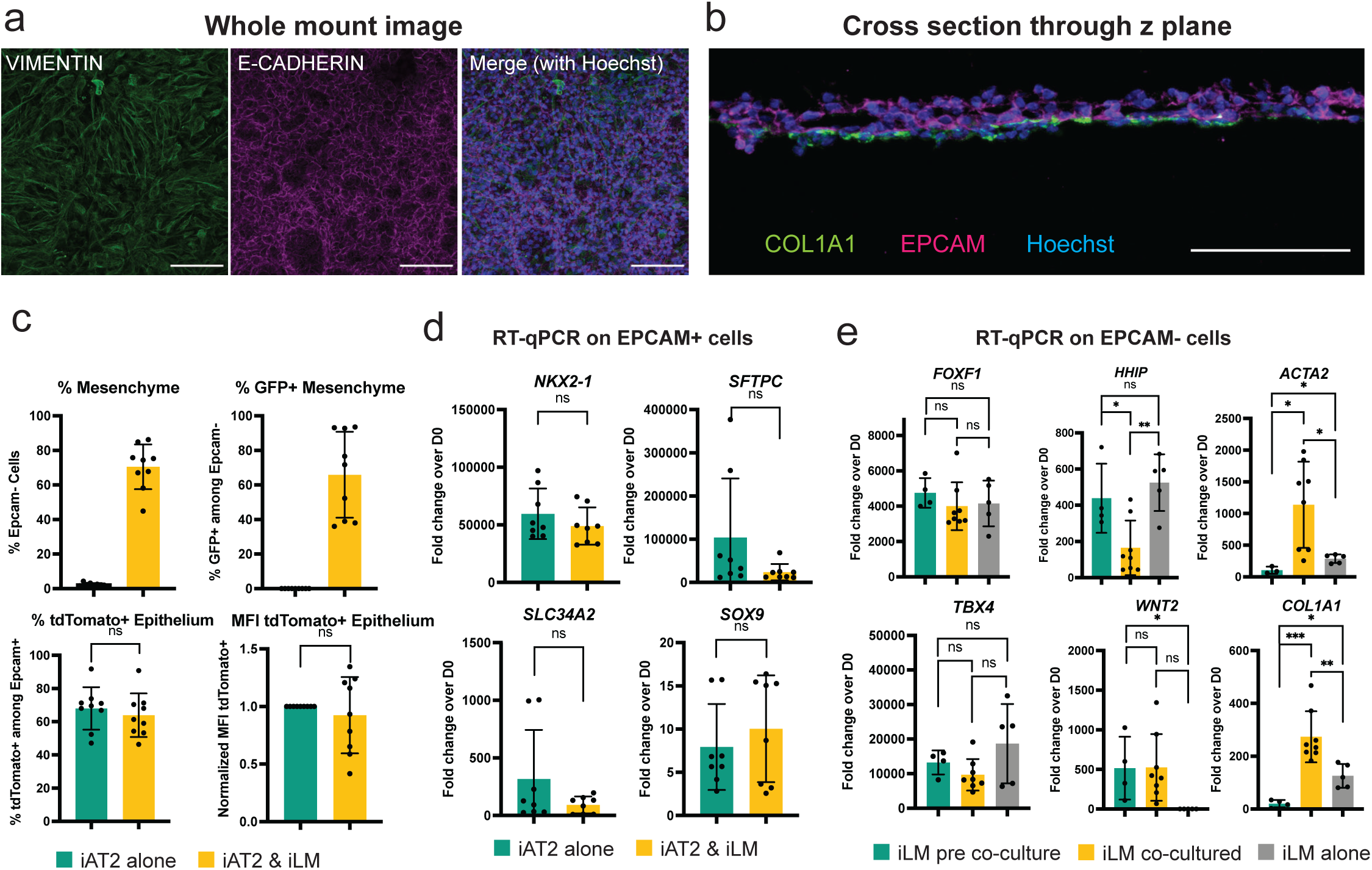
Establishing co-cultures of iLM and iAT2s. **a**: Representative confocal fluorescence microscopy image of a 2D submerged co-culture fixed and immunostained for VIMENTIN and E-CADHERIN. Image represents a maximum intensity z projection from multiple z planes acquired from the bottom to the top of the mesenchymal and epithelial cell layers. Scale bars = 100 μm. Hoechst = nuclei. **b**: Immunofluorescence staining showing expression of COL1A1 and EPCAM in a cross-section of 2D submerged iLM/iAT2 co-cultures. Scale bar = 100 μm. Hoechst = nuclei. **c**: Percentages of EPCAM- cells, of GFP+ cells among EPCAM- cells, and of tdTomato+ cells among EPCAM+ cells, as well as mean fluorescence intensity (MFI) of EPCAM+/tdTomato+ cells after 2D co-culture or in iAT2s cultured alone in the same conditions. N = 9, bars represent mean ± SD, * p<0.05, ** p<0.01, *** p<0.001, ns = not significant, as determined by unpaired two-tailed student’s t test. **d**: Fold change expression over day 0 iPSCs of lung epithelial markers of interest in EPCAM+ sorted cells after 2D co-culture or in iAT2 cells cultured alone in the same conditions. N = 8, bars represent mean and standard deviation. * p<0.05, ** p<0.01, *** p<0.001, as determined by unpaired two-tailed student’s t test. **e**: Fold change expression over day 0 iPSCs of lung mesenchyme markers of interest in iLM cells before co-culture, EPCAM- sorted cells after 2D co-culture, and iLM cultured alone in the same conditions. N = 4 for iLM before co-culture, N = 8 for co-cultured iLM and N = 5 for iLM cultured alone. Bars represent mean ± SD, * p<0.05, ** p<0.01, *** p<0.001, ns = not significant, as determined by unpaired two-tailed student’s t test.

**Supplemental Figure 5:**
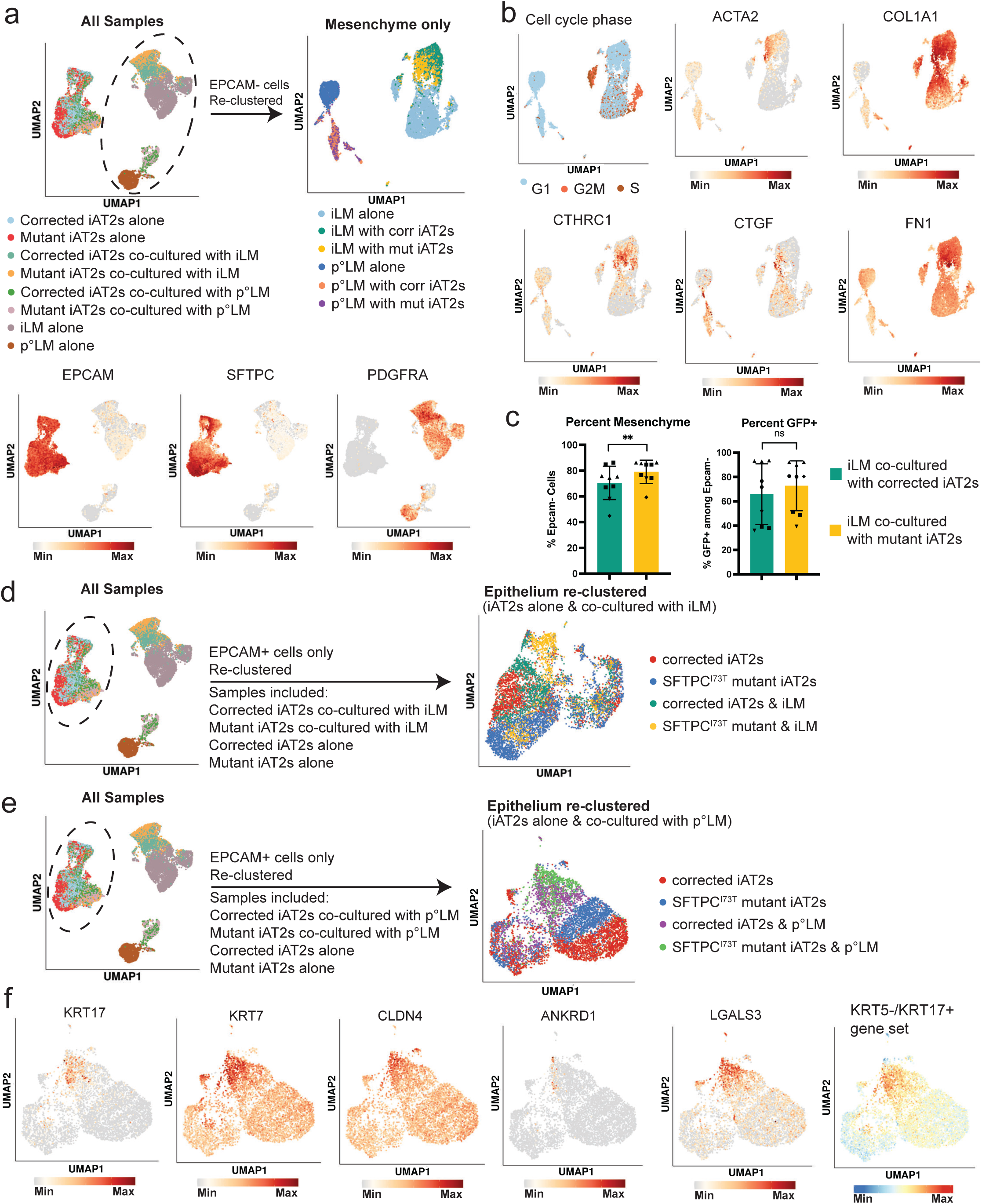
ScRNAseq of co-cultures of iLM with SFTPC^I73T^ mutant vs. corrected iAT2s. **a**: Overview of all samples in the scRNAseq dataset, including co-cultures of iLM or primary lung mesenchyme with SFTPC^I73T^ mutant or corrected iAT2s, as well as single lineage culture controls. UMAP plots show expression of *EPCAM*, *SFTPC* and *PDGFRA* to indicate the epithelial and mesenchymal clusters. “Mesenchyme only” UMAP shows sub-clustering of all EPCAM- clusters. **b**: UMAP plots showing cell cycle phase and expression of fibrotic markers *ACTA2*, *COL1A1*, *CTHRC1*, *CTGF* and *FN1* in re-clustered EPCAM- populations. **c**: Percentage of EPCAM- cells and percentage of GFP+ cells among EPCAM- cells after co-culture of iLM with SFTPC^I73T^ mutant corrected iAT2s for 1 week, as quantified by flow cytometry. N = 8, symbol shapes indicate experimental replicates, i.e. multiple wells of the same experiment, representative of results from 5 repeated independent experiments performed with iAT2s of varying ages and passages. Bars represent mean ± SD, * p<0.05, ** p<0.01, *** p<0.001, ns = not significant, as determined by by paired two-tailed student’s t test. **d**: Overview of all samples in the scRNAseq dataset, as well as subclustered UMAP showing only EPCAM+ clusters from the following samples: SFTPC^I73T^ mutant iAT2s co-cultured with iLM, SFTPC^I73T^ corrected iAT2s co-cultured with iLM, SFTPC^I73T^ mutant iAT2s cultured alone, and SFTPC^I73T^ corrected iAT2s cultured alone. **e**: Overview of all samples in the scRNAseq dataset, as well as subclustered UMAP containing only EPCAM+ clusters from the following samples: SFTPC^I73T^ mutant iAT2s co-cultured with primary human fetal lung mesenchyme, SFTPC^I73T^ corrected iAT2s co-cultured with primary human fetal lung mesenchyme, SFTPC^I73T^ mutant iAT2s cultured alone, and SFTPC^I73T^ corrected iAT2s cultured alone. **f:** UMAP representation of EPCAM+ clusters of SFTPC^I73T^ mutant and corrected iAT2s cultured alone or co-cultured with primary human fetal lung mesenchyme, showing expression of ABI cell markers of interest, as well as expression of KRT5-/KRT17+ gene set from Habermann et al^15^.

**Supplemental Figure 6:**
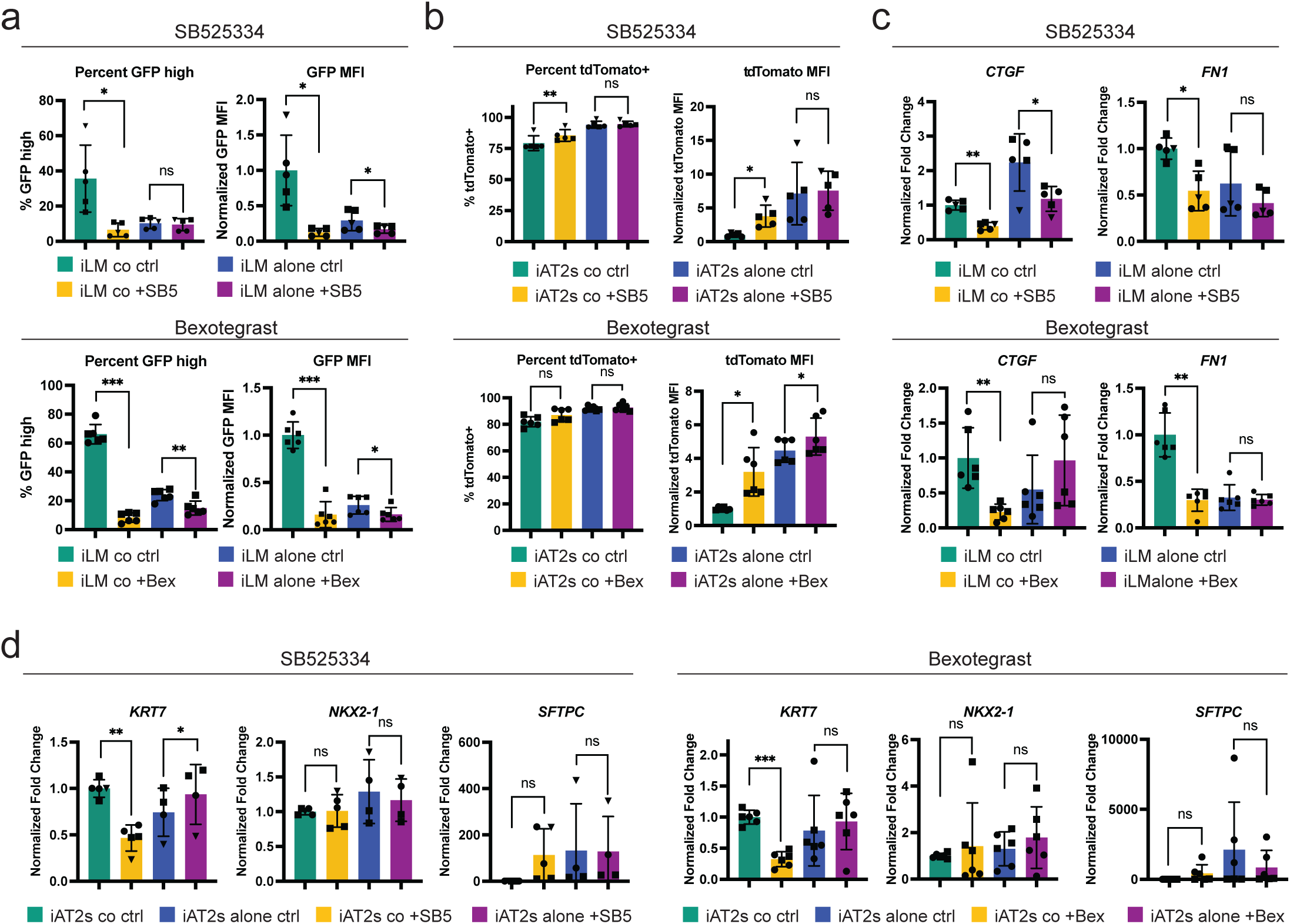
Treatment of co-cultures with SB525334 or Bexotegrast decreases fibrotic activation of the mesenchyme and expression of ABI markers in the epithelium. **a:** Percentage of ACTA2^GFP^ high cells among EPCAM- cells, and mean ACTA2^GFP^ fluorescence intensity of EPCAM- cells in iLM co-cultured with SFTPC^I73T^ mutant iAT2s or cultured alone, treated with 1 μM SB525334 or 2 μM Bexotegrast for 1 week, as compared to vehicle control (treatment starting on the first day of co-culture). N = 5 for SB525334, N = 6 for Bexotegrast, symbol shapes indicate experimental replicates, i.e. multiple wells of the same experiment. Data points are representative of results from 3 repeated independent experiments for SB525334 and 2 repeated independent experiments for Bexotegrast, performed with iAT2s of varying ages and passages. Bars represent mean ± SD, * p<0.05, ** p<0.01, *** p<0.001, ns = not significant, as determined by paired two-tailed Student’s t test. SB5 = SB525334, Bex = Bexotegrast, ctrl = vehicle control, co = co-cultured, alone = cultured alone. **b:** Percentage of tdTomato+ cells among EPCAM+ cells and normalized mean tdTomato fluorescence intensity of EPCAM+/tdTomato+ cells in SFTPC^I73T^ mutant iAT2s co-cultured with iLM or cultured alone, treated as in (a). N = 5 for SB525334, N = 6 for Bexotegrast, symbol shapes indicate experimental replicates, i.e. multiple wells of the same experiment. Data points are representative of results from 3 repeated independent experiments for SB525334 and 2 repeated independent experiments for Bexotegrast, performed with iAT2s of varying ages and passages Bars represent mean ± SD, * p<0.05, ** p<0.01, *** p<0.001, ns = not significant, as determined by paired two-tailed Student’s t test. **c:** RT-qPCR analysis showing fold change expression (normalized to iLM co-cultured control) of fibrotic markers in EPCAM- sorted cells from co-cultures of iLM with SFTPC^I73T^ mutant iAT2s or iLM cultured alone, and treated as in (a). N = 5 for SB525334, N = 6 for Bexotegrast, symbol shapes indicate experimental replicates, i.e. multiple wells of the same experiment. Data points are representative of results from 3 repeated independent experiments for SB525334 and 2 repeated independent experiments for Bexotegrast, performed with iAT2s of varying ages and passages. Bars represent mean ± SD, * p<0.05, ** p<0.01, *** p<0.001, ns = not significant, as determined by paired two-tailed Student’s t test. **d:** RT-qPCR analysis showing fold change expression (normalized to iAT2 co-cultured control) of *KRT7, NKX2-1* and *SFTPC* in EPCAM+ sorted cells from SFTPC^I73T^ mutant iAT2s co-cultured with iLM or cultured alone, and treated as in (a). N = 5 for SB525334, N = 6 for Bexotegrast, symbol shapes indicate experimental replicates, i.e. multiple wells of the same experiment. Data points are representative of results from 3 repeated independent experiments for SB525334 and 2 repeated independent experiments for Bexotegrast, performed with iAT2s of varying ages and passages. Bars represent mean ± SD, * p<0.05, ** p<0.01, *** p<0.001, ns = not significant, as determined by paired two-tailed Student’s t test.

## Methods

### Cell lines

The BU3 ACTA2^GFP^/TBX4-LER^tdTomato^ (BU3 A2GTT) reporter iPSC line was generated by adding a TBX4 lung enhancer reporter construct to our previously published BU3 ACTA2^GFP^ reporter iPSC line^76^. The TBX4-LER^tdTomato^ reporter construct was engineered into the BU3 ACTA2^GFP^ reporter iPSC line using zinc finger nucleases (ZFNs) targeting the AAVS1 locus. In brief, ZFNs were used to induce a double stranded break at the AAVS1 locus, and a donor plasmid with left and right homology arms to the AAVS1 locus target site flanking a splice acceptor, T2A peptide, and antibiotic selection cassette, as well as the TBX4 lung enhancer sequence, minimal CMV promoter, and tdTomato fluorescent reporter was provided for targeted integration by homologous recombination. Both ZFNs and donor plasmids were introduced into iPSCs by nucleofection using the Lonza P3 Primary Cell 4D Nucleofector^TM^ X Kit (Lonza, #V4XP-3024). Puromycin-resistant clones were mechanically isolated and plated into fresh Matrigel-coated plates for clonal expansion. Targeted cells were confirmed by PCR screening using primers binding outside of the 5’ homology arm and within the antibiotic resistance cassette (Supplemental Figure 1a). In addition, primers within the homology arms were used to screen for mono-allelic or bi-allelic targeting of the ACTA2 locus (Supplemental Figure 1a).

All iPSC lines used in this study, both before and after gene editing, had a normal karyotype when analyzed by G-banding (Cell Line Genetics) and pluripotency was confirmed by staining for pluripotency markers as previously described^98^. Prior to directed differentiation, iPSC lines were maintained in feeder-free conditions on growth factor reduced 2D Matrigel (Corning, #354277) in 6-well tissue culture plates. The BU3 A2GTT iPSC line was cultured in StemFlex medium (Thermo Fisher Scientific, #A3349401), the SFTPC^tdT/WT^ (aka “SFTPC corrected”)^51^ and SFTPC^I73T/tdT^ (aka “SFTPC^73T^ mutant”)^51^, BU1^99^, and RUES2^100^ (generous gift from A. Brivanlou, Rockefeller University) iPSC/ESC lines were cultured in mTeSR1 medium (StemCell Technologies, #05850). Gentle cell dissociation reagent (StemCell Technologies, #07174) was used for passaging. All iPSC/ESC differentiations were performed under regulatory approval of the Institutional Review Board of Boston University. All iPSC lines, including BU3 A2GTT, can be requested for sharing through www.kottonlab.com.

### Primary developing bud tip epithelium and mesenchyme

Human lung tissue research involving the use of human developing bud tip epithelium and mesenchyme was reviewed and approved by The University of Michigan Institutional Review Board (IRB). Normal, de-identified human lung tissue was obtained from the University of Washington Laboratory of Developmental Biology. Tissue was shipped overnight in UW-Belzer’s solution (Thermo Fisher, #NC0952695) on ice and was processed for experiments within 24h. Bud tip epithelial progenitors (BTE) were isolated from tissue from day 132 post conception and cultured as previously described^101^. In brief, lung specimens were minced into small 2 mm x 2 mm pieces, washed with 1 x PBS and then treated for 45 min with Dispase (Gibco, #17105-041) at a concentration of 50 caseinolytic units per ml in order to dissociate mesenchyme while keeping the epithelium intact. Dispase was quenched with 100 % fetal bovine serum (FBS, Corning, #35015CV) for 10 min and then pipetted vigorously in DMEM/F12 (Corning, #10092CV) with 10 % FBS (Gibco, #A5670801) until intact epithelial bud tips were separated from dissociated mesenchymal cells. BTE organoid medium consisted of DMEM/F-12, 100 U/mL penicillin-streptomycin (Thermo Fisher, #15140122), 1 × B-27 supplement (Invitrogen, #17504044), 1 × N2 supplement (Invitrogen, #17502048), 0.05 % BSA (Sigma, #A9647), 50 μg/mL L-ascorbic acid (Sigma, #A4544), 0.4 μM 1-Thioglycerol (Sigma, M1753), 50 nM all-trans retinoic acid (Sigma, #R2625), 10 ng/mL recombinant human FGF7 (R&D Systems, #251-KG) and 3 μM CHIR99021 (APExBIO, Cat#A3011), and was replenished every 3 days. After isolation, approximately 100 intact bud tips were seeded in 40 µl droplets of Matrigel (Corning, #354234) diluted to 8 mg/mL before use. BTE organoids were passaged at a ratio of 1/4 – 1/6 by needle sheering every 7 – 10 days. For co-culture, BTE was collected from Matrigel by pipette and dissociated by needle sheering and combined with dissociated TBX4-LER^tdTomato^+ cells such that BTE Organoids underwent a 1:4 split and then were reseeded in 40 µl droplets of Matrigel containing 7000 TBX4-LER^tdTomato^+ cells per µl. Recombinant cultures were maintained in BTE Organoid medium without CHIR99021. For the first 24 h 10 µM Y-27632 (Fisher #125450) was included in the media. Media was exchanged at least every 3 days. After 10 days, organoids were collected and processed for immunofluorescence and qPCR.

Primary lung mesenchyme from day 57, 76, 94 and 122 post conception was collected from completed bud tip progenitor organoid isolations. Mesenchymal cells were further enriched by centrifugation at 10xG for 1 min at 4 °C, which pellets intact epithelium while leaving dissociated mesenchymal cells in suspension. Suspension was collected and cells were counted by hemocytometer and cryopreserved in 1 mL of DMEM/F12 with 10 % Dimethyl Sulfoxide (Sigma #D8418) and 10 % FBS at a concentration of 1 – 3×10^6^ cells per vial.

### Primary adult lung fibroblasts

Human primary adult lung fibroblasts were isolated from distal lung tissue from a de-identified healthy donor (66y, female, white, non-smoker, non-alcoholic) procured from the International Institute for the Advancement of Medicine (IIAM). The distal tissue was cut into 1 cm × 1 cm pieces and kept in 6-well plates submerged in Advanced DMEM/F-12 (Gibco, #12634010), with 10 % FBS (Gibco, #A5670801), 1 % non-essential amino acids (Sigma-Aldrich, #M7145) and 1 % Glutamax (Gibco, #36050-061) for 4 weeks to allow the fibroblasts to dislodge from the tissue and form an adherent monolayer in the well. The adherent cell population was then dissociated with TrypLE (Gibco, #12605036), cultured in cell culture flasks and passaged once a week until co-culture (passage 7).

### Directed differentiation of human iPSCs into the lung mesenchyme

For directed differentiation of iPSCs into lung mesenchyme, cells were first differentiated into lateral plate mesoderm as previously published^76^. In brief, cells were dissociated using Gentle Cell Dissociation Reagent (StemCell Technologies, #07174) and 1×10^6^ cells were plated onto a well of a 6-well tissue culture plate coated with growth factor reduced 2D Matrigel (Corning, #354277) in StemFlex or mTeSR1 medium supplemented with 10 mM Y-27632 (Tocris, #1254). After 24 h (i.e. on Day 0 of differentiation) iPSCs were treated with Stemline II Hematopoietic Stem Cell Expansion medium (Sigma-Aldrich, #S0192) supplemented with 10 ng/mL each of rhBMP4 (R&D Systems, #314-BP), rhActivin A (R&D Systems, #338-AC), rhVEGF (R&D Systems, #293-VE), and rhFGF2 (R&D Systems, #233-FB) for 24 h, followed by 72 h of treatment with 10 ng/mL each of rhBMP4, rhVEGF, and rhFGF2. On day 4 of differentiation, differentiation efficiency was assessed by flow cytometry for KDR expression using a mouse anti-human CD309 antibody (BD Biosciences, #560494, 1/50). Cells were dissociated into small clumps using Gentle Cell Dissociation Reagent and passaged at 1/50 onto 12-well tissue culture plates coated with growth factor reduced Matrigel (Corning 354277) in complete serum-free differentiation medium (cSFDM) supplemented with 2 μM Retinoic Acid (Sigma, #R2625), 1 μM SAG (StemCell Technologies, #73412), 1 μM CHIR99021 (Fisher, #44-235-0) and 10 mM Y-27632 (Tocris, #1254). 500 ml cSFDM was composed of 375 ml IMDM (ThermoFisher, #12440053), 125 ml Ham’s F12 (Cellgro, #10-080-CV), 5 ml Glutamax 100x (Gibco, #35050-061), 5 ml B27 supplement without vitamin A (Invitrogen, #12587-010), 2.5 ml N2 (Invitrogen, #17502-048), 3.3 ml 7.5 % BSA (Life Technologies, #15260-037), 19.5 μL 1-thioglycerol(final concentration 4.5 × 10-4 M, Sigma, #M6145), 500 μl ascorbic acid (final concentration 50 μg/ml, Sigma, #A4544), and 1 ml Primocin (final concentration 100 μg/ml, Fisher Scientific, #NC9392943). After 48 h medium was replaced with fresh RCS medium (i.e. cSFDM medium containing 2 μM Retinoic Acid, 1 μM SAG, and 1 μM CHIR99021) without Y-27632 and the medium was replaced every 48 h. On day 16 of differentiation, cells were passaged using 0.05 % Trypsin (Gibco, #25300062) for 4 min at 37 °C and subsequent inactivated with Fetal Bovine Serum. Cells were centrifuged for 5 min at 300xG, resuspended in fresh RCS medium and re-plated at 1/12 onto fresh Matrigel-coated 12-well tissue culture plates in RCS medium with 10 mM Y-27632. RCS medium was replaced every 48 h until collection. Cells were collected for further analysis between day 16 and day 30 of differentiation as indicated in the text, using 0.05 % Trypsin for 4 min at 37 °C and subsequent inactivation with Fetal Bovine Serum. TBX4-LER^tdTomato^ percentage was quantified using a Stratedigm flow cytometer, and cell sorting was performed at the Boston University Flow Cytometry Core Facility using a MoFlo Astrios cell sorter.

### Directed differentiation of iPSCs into alveolar epithelial type 2 cells (iAT2s) and iAT2 cell maintenance

We used our previously published SFTPC^tdT/WT^ (“SFTPC corrected”) and SFTPC^I73T/tdT^ (“SFTPC^73T^ mutant”) iPSC lines^51^ and our previously published directed differentiation protocol^50,51^ in order to generate SFTPC^I73T^ mutant and corrected iAT2s. In brief, the STEMdiff Definitive Endoderm Kit (Stemcell Technologies, #05110) was used for differentiation into endoderm. On day 3, cells were dissociated using Gentle Cell Dissociation Reagent (StemCell Technologies, #07174), passaged onto 6 well tissue culture plates coated with growth factor reduced Matrigel (Corning 354277) and cultured in cSFDM supplemented with 2 μM Dorsomorphin (“DS”, Fisher Scientific, #NC0275327) and 10 μM SB431542 (“SB”, Sigma, #S4317). Note that that the composition of cSFDM medium is the same as for the iLM differentiation described above, with the exception of the B27 supplement. For all iAT2 differentiations and cultures, as well as all iLM/iAT2 co-culture experiments B27 with vitamin A was used (Invitrogen, # 17504-44). After 3 days of culture in “DS/SB” medium cells were cultured in cSFDM medium supplemented with 3 μM CHIR99021, 10 ng/ml rhBMP4 (R&D Systems, #314-BP) and 100 nM Retinoic Acid (Sigma, #R2625). On day 15-16 of differentiation cells were sorted for lung progenitors based on CD47^hi^/CD26^neg^ gating. Sorted cells were resuspended in 50 µl droplets of 3D Matrigel (Corning, #354230) at 400 cells/μl and cultured in alveolar differentiation medium “CK+DCI”, consisting of cSFDM supplemented with 3 μM CHIR99021 (Fisher Scientific, 44-235-0), 10 ng/ml rhKGF (Fisher Scientific, #251KG050), 50 nM Dexamethasone (Sigma, #D4902), 0.1 mM 8-bromoadenosine 3’,5’ cyclic monophosphate sodium salt (Sigma, # B7880) and 0.1 mM 3-isobutyl-1-methylxanthine (IBMX, Sigma, # I5879). CK+DCI medium was supplemented with 10 mM Y-27632 (Tocris, #1254) for the first 48 h after sorting, and cells were fed with fresh CK+DCI medium every 48-72 h. Epithelial spheres were passaged between day 28 and 37 of differentiation, and a brief period of CHIR withdrawal was subsequently performed to achieve iAT2 maturation as previously described^43^. SFTPC^tdTomato^+ cells were then purified by FACS sorting and plated at 400 cells/μl in 3D Matrigel (Corning, #354230). iAT2s were subsequently maintained through serial passaging and re-plating into 3D Matrigel droplets at a density of 400 cells/μl every 10 days. For passaging, cells were dissociated into single cells with 2 mg/ml Dispase (Gibco, #17105-041) for 1 h at 37 °C, followed by incubation in 0.05 % Trypsin (Gibco, #25300062) at 37 °C for 12 min. iAT2 cell purity was assessed by flow cytometry for SFTPC^tdTomato^ expression. iAT2s between day 100 – 200 of differentiation were used for co-culture with iLM cells.

### Co-culture of human iLM and iAT2s

For 2D co-culture, TBX4-LER^tdTomato^+ cells were sorted on day 26 – 30 of differentiation and plated onto 96-well tissue culture plates or, if processed for immunofluorescence, 24-well trans-well inserts (Corning, #3470) coated with fresh 2D Matrigel (Corning, #354277) at 150’000 cells per well in RCS medium with 10 mM Y-27632. For culture on 96-well plates 200 μl of RCS medium was used, for culture on trans-well inserts 100 μl of medium was added to the apical compartment and 600 μl of medium to the basolateral compartment. After 48 h, iAT2s were dissociated from 3D Matrigel droplets as described above and subsequently plated on top of iLM cells at 250’000 cells per well in CK+DCI medium with 10 mM Y-27632 (200 μl of medium per well in 96-well plates, 100 μl of medium in the apical compartment and 600 μl of medium in the basolateral compartment for trans-well inserts). Medium was replaced with fresh CK+DCI medium every 48 h. For SB525334 or Bexotegrast treatment of co-cultures, 1 μM SB525334 (Sigma, #S8822-5MG) or 2 μM Bexotegrast (MedChemExpress, #HY-137561) was added to the co-cultures at the time-point of iAT2 plating and kept in the culture medium until the time-point of cell collection.

Cells were harvested after 7 days of co-culture. To re-isolate iAT2 and iLM cells after 2D co-culture, cells were incubated in 200 μl Accutase (Sigma Aldrich, #A6964) for 20 min at room temperature. Cells were pipetted up and down several times every 5 min to dislodge clumps and then placed at 37 °C for 15 min additional min before cell collection and centrifugation at 300xG for 5 min. Dissociated cells were incubated with an anti-EPCAM antibody (BD Biosciences, #563180, 1/500) for 30 min on ice. Cells were collected by centrifugation at 300xG for 5 min and resuspended in FACS buffer (PBS with 2 % FBS) supplemented with the live/dead stain DRAQ7 (Biolegend, #424001, 1/100). For sorting, cells were gated for single cells and live cells, followed by separation of EPCAM+ and EPCAM- cells. All cell sorting was performed at the Boston University Flow Cytometry Core Facility using a MoFlo Astrios sorter.

### Reverse transcriptase quantitative real time polymerase chain reaction (RT-qPCR)

For RT-qPCR of iLM co-cultured with bud tip epithelium (BTE), organoids were manually dislodged from Matrigel using a P1000 pipette tip, transferred into 1.5 mL microcentrifuge tubes, and flash-frozen by immersing the tubes in a small volume of liquid nitrogen, ensuring minimal residual culture media. Total RNA was extracted from the frozen pellets using the MagMax-96 Total RNA Isolation Kit (Thermo Fisher, #AM1830), following the manufacturer’s protocol. RNA concentration and purity were assessed using a NanoDrop 2000 spectrophotometer. Reverse transcription was carried out in technical triplicates using the SuperScript VILO cDNA Synthesis Kit, with 200 ng of input RNA per reaction. The resulting cDNA was diluted 1/2 with DNase/RNase-free water, and 1/40th of the total reaction volume was used for each quantitative PCR reaction. RT-qPCR was performed on a StepOnePlus Real-Time PCR System (Thermo Fisher, #4376559R) using QuantiTect SYBR Green qPCR Master Mix (Qiagen, Cat#204145), with primers (Supplemental Table 9) used at a final concentration of 500 nM. Arbitrary units of gene expression were calculated using 18S rRNA as a normalizer using the following equation: Arbitrary Units (AUs) = 2^(18SCt - GeneCt)^ ✕ 10,000.

For all other RT-qPCR data shown in this manuscript cells were collected by FACS. For iLM differentiations cells were sorted based on TBX4-LER^tdTomato^ expression. For RT-qPCR of co-cultures, mesenchyme and epithelium were re-separated and sorted based on EPCAM expression as described above. Sorted cells were centrifuged at 300xG for 5 min. Supernatant was removed and cell pellets were resuspended in 700 μl QIAzol Lysis reagent (Qiagen, #79306). RNA was extracted using a RNeasy Plus Mini Kit (QIAGEN, #74136) as instructed by the manufacturer. RNA was eluted in 30 μl nuclease-free water and RNA concentration was measured using a NanoDrop ND-1000 microvolume spectrophotometer (Thermo Fisher Scientific, Waltham, MA). cDNA was generated from 150 ng of RNA in a 20 μl reaction using a High-Capacity cDNA Reverse Transcription Kit (Applied Biosystems, #4368814) and cDNA was diluted 1/16 in nuclease-free water. qPCR was performed using TaqMan Fast Universal PCR Master Mix (Thermo Fisher, #364103), using 4 μl of diluted cDNA in a 10 μl qPCR reaction. qPCR was run on an Applied Biosystems QuantStudio7 384-well system with a cycle number of 40. Relative expression was calculated using the the 2^(−ΔΔCt)^ method^102^, using 18S rRNA (Thermo Fisher, #4318839) as the internal reference gene. All reactions were run in technical duplicates. Undetected Ct values were set to the maximum number of cycles (40) to allow fold change calculations. TaqMan gene expression arrays were purchased from Thermo Fisher.

### Immunofluorescence microscopy

Widefield images of live cultures were acquired using a Keyence BZ-X700 fluorescence microscope. Confocal microscopy was performed at the Boston University Cellular Imaging Core, using a Zeiss LSM710 microscope.

For immunofluorescence imaging of day 28 and 30 iLM cells (Figure 1d, Supplemental Figure 2a), cells were plated into gelatin-coated glass chamber slides (Millipore, #PEZGS0816) on day 16 and cultured until day 28/30 in RCS medium. Cells were then fixed with 4 % paraformaldehyde for 10 min at room temperature, permeabilized with 0.25 % Triton X-100 and 2.5 % normal donkey serum for 30 min at room temperature, and subsequently blocked with 2.5% normal donkey serum for 20 min at room temperature. Cells were incubated with primary antibody (rabbit anti-RFP, Rockland #600-401-379, 1/250; chicken anti-GFP, Aves Labs #GFP-1020, 1/500; goat anti-FOXF1, R&D Systems #AF4798, 1/500) in 5 % normal donkey serum over night at 4 °C. Cells were washed 3 times with PBS, incubated with secondary antibody and Hoechst 33342 (Thermo Fisher Scientific, # H3570, 1/500) for 2 h at room temperature, mounted with Prolong Diamond Antifade Reagent (Invitrogen, # P36965) and imaged with a Zeiss LSM710 confocal microscope.

For immunofluorescence imaging of iLM co-cultured with primary bud tip epithelium (BTE), samples were fixed for 24 h in 10 % Neutral Buffered Formalin at room temperature with gentle agitation. Samples were washed 3 × with distilled water and dehydrated through washes in 25 %, 50 %, 75 %, and 100 % methanol. Samples were then transferred to 100 % ethanol followed by 70 % ethanol and embedded in Histogel (VWR, #73009-992). After fixation, each step was performed for at least 15 min at room temperature with gentle agitation. Processing for paraffin embedding occurred in an automated tissue processor with 1 h each step through the following series of solutions: 70 %, 80 %, 2 × 95 %, 3 × 100 % ethanol, 3 × xylene, and 3 × paraffin. Histogel embedded paraffin processed samples were embedded into paraffin blocks and cut into 5 μm-thick sections onto charged glass slides using a microtome. Slides were baked for 1 h at 60 °C immediately prior to staining. Slides then underwent two 5 min washes with HistoClear II (National Diagnostics, #HS-202), followed by a graded ethanol rehydration series (100 %, 95 %, 70 %, and 30 %) for 6 min per step, with buffer replacement midway through each incubation. Following rehydration, slides were rinsed twice in double-distilled water (ddH₂O) for 5 min each. Antigen retrieval was carried out using 1× Sodium Citrate Buffer (comprising 100 mM trisodium citrate (Sigma, #S1804) and 0.5 % Tween-20 (Thermo Fisher, #BP337, pH 6.0) in a steamer under atmospheric pressure for 20 min. After three washes in ddH₂O, slides were blocked for 1 h using a solution containing 5 % normal donkey serum (Sigma, #D9663) and 0.1 % Tween-20 in PBS. Primary antibodies (rabbit anti TAGLN, Abcam #ab14106, 1/500; mouse anti-CDH1, BD Biosciences #610182, 1/500; goat anti-VIMENTIN, R&D Systems #AF2105, 1/500; rabbit anti-TP63, Cell Signaling Technologies #13109, 1/500; mouse anti-SPC, Santa Cruz Biotechnology #SC-518029, 1/200; rabbit anti-SOX9, Millipore #AB5535, 1/500; goat anti-SOX2, R&D Systems #AF2018, 1/500), diluted in the same blocking buffer, were applied and incubated overnight at 4 °C in a humidified chamber. The next day, slides were washed three times with 1× PBS for 10 min each. Secondary antibodies and DAPI (1 μg/mL), also diluted in blocking buffer, were applied for 1 h, followed by three 5-min washes in PBS. Finally, slides were mounted with ProLong Gold (Invitrogen, #P369300) and imaged within two weeks.

For immunofluorescence imaging of co-cultures, co-cultures were plated on 24-well trans-well inserts (Corning, #3470). After 7 days of co-culture, cells were fixed with 4 % paraformaldehyde for 15 min at room temperature, permeabilized with 0.1 % Triton X-100 for 10 min and blocked with 4 % normal donkey serum and 0.1 % Triton X-100 for 1 h at room temperature. Trans-well inserts were then incubated with primary antibody (mouse anti-VIMENTIN, Thermo Fisher #MA5-11883; 1/100; rabbit anti-ECADHERIN, Cell Signaling Technology #3195S, 1/1600; rabbit anti-Cytokeratin 17 (KRT17)-Alexa Fluor 488, Abcam, #ab185032; rabbit anti-Cytokeratin 17 (KRT17), Abcam, #ab109725) in 4 % normal donkey serum at 4 °C overnight, washed 3 x with PBS and incubated with secondary antibody and Hoechst 33342 (Thermo Fisher Scientific, # H3570, 1/500) for 1 h at room temperature. Trans-well insert membranes were cut out with a razor blade and mounted onto slides with ProLong Diamond Antifade Mountant (Invitrogen, #P36965).

For cross-section imaging of co-cultures, cells were cultured on 24-well trans-well inserts (Corning, #3470), fixed with 4 % paraformaldehyde for 10 min at room temperature and washed 3 x with PBS. Membranes were incubated with 30 % sucrose in PBS at 4 °C over night. Membranes were then cut out using a razor blade and embedded in plastic molds filled with O.C.T. compound (Fisher Scientific, # 23-730-571). Molds were stored on dry ice for 10-20 min to let O.C.T. solidify and subsequently stored at −80° C. O.C.T. blocks were sectioned at 12 μm per slice and transferred onto slides. For immunofluorescence, slides were washed 3 x with PBS, blocked with 4 % Normal donkey serum and 0.1 % Triton X for 1 h and subsequently incubated with primary antibody (mouse anti-VIMENTIN, Thermo Fisher #MA5-11883, 1/100; rabbit anti-ECADHERIN, Cell Signaling Technology #3195S, 1/1600; rabbit anti-SMA, Abcam #ab5694, 1/200; mouse anti-EPCAM, Abcam #ab20160, 1/200; rabbit anti-COL1A1, Cell Signaling Technology #72026T, 1/150), and secondary antibody and Hoechst 33342 (Thermo Fisher Scientific, #H3570, 1/500) as described above. Slides were imaged with a Zeiss LSM710 confocal microscope.

For the Picrosirius red stains, a stain kit was used (Abcam, #ab150681) and staining was performed as instructed by the manufacturer. Staining was imaged using a Keyence light microscope using the iLM only control to set the white balance.

### Single cell RNA sequencing (scRNAseq)

For single cell RNA sequencing, cells were differentiated as described and sorted based on expression of fluorescent markers as indicated in the text. Primary developing lung mesenchyme cells were thawed and sorted for live cells using Calcein Blue (Life Technologies, #C1429). Single cell capture (10× Chromium instrument; 10× Genomics, Pleasanton, CA), library preparation (Chromium Single-Cell 30 Library Kit; 10× Genomics), and sequencing was performed at the Boston University Single Cell Sequencing Core Facility. Sequencing libraries were loaded on an Illumina NextSeq2000 with a custom sequencing setting to obtain a sequencing depth of ∼40k reads/cell. Sequencing files were mapped to the human genome reference (GRCh37) supplemented with GFP and tdTomato sequences using CellRanger v3.0.2. Seurat v3.2.3 for downstream analysis and quality control^103^. After inspection of the quality control metrics, cells with 15 % to 35 % of mitochondrial content and <800 detected genes were excluded for downstream analyses. In addition, doublets were also excluded for downstream analysis. We normalized and scaled the unique molecular identifier (UMI) counts using the regularized negative binomial regression (SCTransform)^104^. Following the standard procedure in Seurat’s pipeline, we performed linear dimensionality reduction (principal component analysis) and used the top 20 principal components to compute the unsupervised Uniform Manifold Approximation and Projection (UMAP)^105^. For clustering of the cells, we used Louvain algorithm which were computed at a range of resolutions from 1.5 to 0.05 (more to fewer clusters)^106^. Cell cycle scores and classifications were done using the Seurat’s cell-cycle scoring and regression method ^107^. Cluster-specific or sample-specific genes were calculated using MAST framework in Seurat^108^. An online Shiny app has been established to allow interactive, user-friendly visualizations of gene expression in each population^109^.

### Differential ligand-receptor interaction analysis using NicheNet

To infer ligand–receptor interactions underlying epithelial–mesenchymal crosstalk, we performed differential NicheNet analysis using the NicheNetR package version 2.2.0 in R^86^, following the workflow provided in the NicheNet differential analysis vignette (https://github.com/saeyslab/nichenetr/blob/master/vignettes/differential_nichenet.md). scRNAseq data were pre-processed and annotated using Seurat (v3.2.3). Sender and receiver cell populations were defined as iAT2s and iLM, or iLM and iAT2s respectively, with analyses performed on the following conditions: (1) iLM cultured alone, (2) iLM co-cultured with SFTPC^I73T^ mutant iAT2s, (3) iLM co-cultured with corrected iAT2s, (4) SFTPC^I73T^ mutant iAT2s cultured alone, (5) corrected iAT2s cultured alone, (6) SFTPC^I73T^ mutant iAT2s co-cultured with iLM, (7) corrected iAT2s co-cultured with iLM. For each condition, genes expressed in at least 10 % of cells per group were included. Differential expression between conditions was calculated using Seurat’s Wilcoxon rank-sum test. Ligands upregulated in senders and their cognate receptors upregulated in receivers were identified, and ligand activity prediction was performed using the NicheNet regulatory potential scores. Following the NicheNet vignette, we applied the prioritizing weights function to combine multiple features—such as ligand expression in sender cells, receptor expression in receiver cells, and regulatory potential—into a single prioritization score for each ligand–receptor pair. This integrated weighting allowed us to systematically rank and filter pairs, prioritizing those most likely to mediate condition-specific intercellular communication (i.e., those with highest composite scores in fibrotic versus control conditions). Analyses were performed for communication in both directions (iAT2→iLM and iLM→iAT2).

### Single-Cell Type Order Parameters (scTOP) analysis of iLM

Single-Cell Type Order Parameters (scTOP) is a computational method to classify and visualize individual cells by measuring their similarity to a defined set of reference cell types ^79,80^. It provides a quantitative score for cell identity without relying on common analysis techniques such as dimensional reduction, statistical fitting or clustering, and is less susceptible to batch effects than other methods. Previously published single cell atlases can be used to create a reference basis and similarity scores are computed for each cell/population of interest from a scRNAseq dataset of interest as compared to the cell populations in the reference database. For more details we refer to our publication, Yampolskaya et al., 2023^79^. We employed scTOP in order to classify and visualize the similarity of individual cells within our iPSC-derived Day 17 TBX4-LER^tdTomato^+, Day 26 TBX4-LER^tdTomato^+, and Day 26 TBX4-LER^tdTomato^- samples to known mesenchymal cell populations. We used scTOP to rank-order gene counts, z-score cells, and project each cell onto a reference basis normalized by the same procedure and constructed from select populations of a human fetal lung scRNAseq dataset by He et al^77^. Seven mesenchymal populations of interest were included in the reference: early fibroblasts (week 5), adventitial fibroblasts (week 20), alveolar fibroblasts (week 20), myofibroblasts (week 20), airway smooth muscle cells (week 20), pericytes (week 20), and mesothelial cells (week 20). Five non-mesenchymal populations were included as negative controls: early capillary cells (week 5), early airway progenitors (week 5), ciliated cells (week 15), DC1 (week 15), and DC2 (week 15). Note that certain mesenchymal populations from the scRNAseq dataset by He. et al were excluded from the reference basis due to low cell counts or high similarity to other populations, including mesenchymal 1, 2, and 3, and mid fibro. However, we re-introduced these populations into our analysis when building alignment score plots (Figure 2b, Supplemental Figure 3b-c) to compare their projections onto the reference basis to those of our iPSC-derived populations.

### Quantification and statistical analysis

Statistical methods relevant to each figure are outlined in the figure legends. In brief, bar graphs represent mean and standard deviation and each data point represents one biological replicate, i.e. one independent differentiation/co-culture experiment, unless otherwise indicated. In figures that include multiple experimental replicates (i.e. multiple wells of the same differentiation) this is indicated in the figure legends and represented in the figure using different symbol shapes for each biological replicate. Multiple biological replicates (i.e. independent differentiations or co-culture experiments) were used for all experiments, except co-cultures of iLM with primary bud tip epithelium, which were performed as one biological replicate with 3 experimental replicates (i.e. 3 wells of co-cultures plated at the same time). Statistically significant differences between conditions were determined using Student’s t tests or ANOVA, as indicated in the figure legends.

## Acknowledgements

We thank Brian R. Tilton from the Boston University Flow Cytometry Core Facility for cell sorting, and Yuriy Alekseyev and Christopher Williams from the Boston University Single Cell Sequencing Core for scRNA sequencing. Schematics were created with Biorender.com. This work was supported by NIH grants R01HL095993, NO175N92025D00035 and U01HL148692 to DNK, NIH K08 HL163494 and a Boston University School of Medicine Department of Medicine Career Investment Award to KDA, NIH P01HL170952 and U01HL152976 to DNK and KDA, Swiss National Science Foundation P500PB_206631 to ABA, NIH NIGMS 5R35GM119461-09 to PM, HS and MY, NIH R01HL119215 and R01HL166139 to JRS, and National Heart, Lung, and Blood Institute (NHLBI) U01HL153000, NHLBI LungMAP U01HL175451, DoD PR202868 and the Burroughs Wellcome Fund 1020030 to BNG.

## List of Supplemental Tables

**Supplemental Table 1:** List of all DEGs in day 26 TBX4-LER^tdTomato^+ cells, day 26 TBX4-LER^tdTomato^-cells and day 17 TBX4-LER^tdTomato^+ cells.

**Supplemental Table 2:** List of all DEGs in Louvain clusters (resolution 0.5) of day 26 TBX4-LER^Tomato^+ sample.

**Supplemental Table 3:** Gene sets used in Figure 2d. Gene sets were generated from top 50 DEGs in day 57 primary, day 76 primary, day 94 primary and day 122 primary clusters.

**Supplemental Table 4:** List of 80 genes used to generate fibrotic markers gene set shown in Figure 3d. Gene list was generated from Sikkema et al^84^.

**Supplemental Table 5:** List of all DEGs between iLM co-cultured with SFTPC^I73T^ mutant iAT2s vs iLM co-cultured with corrected iAT2s, and all DEGs between primary adult lung mesenchyme co-cultured with SFTPC^I73T^mutant iAT2s vs primary adult lung mesenchyme co-cultured with corrected iAT2s.

**Supplemental Table 6:** List of 20 genes used to generate the KRT5-/KRT17+ gene shown in Figure 4d. Gene list was generated from scRNAseq dataset from Habermann et al^15^.

**Supplemental Table 7:** List of all DEGs between monocultures of SFTPC^I73T^ mutant vs monocultures of corrected iAT2s, and all DEGs between SFTPC^I73T^ mutant vs corrected iAT2s when co-cultured with iLM.

**Supplemental Table 8:** Top 20 prioritized ligands and receptors between co-cultures of iLM with SFTPC^I73T^ mutant iAT2s and co-cultures of iLM with corrected iAT2s, either defining iAT2s as sender and iLM as receiver or vice versa.

**Supplemental Table 9:** List of primers used for RT-qPCR of co-cultures of iLM with primary BTE.

