## Supplementary material for "Bidirectional fibrogenic cross-talk revealed in a human iPSC-derived epithelial-mesenchymal co-culture model of pulmonary fibrosis": Supplememtal Table 9

### Supplemental Table 9: Primers for RT-qPCR on iLM/BTE co-cultures.

| **Primer** | **Sequence (5' → 3')** |
| --- | --- |
| hSOX9_F | GTACCCGCACTTGCACAAC |
| hSOX9_R | GTGGTCCTTCTTGTGCTGC |
| hSOX2_F | TACAGCATGTCCTACTCGCAG |
| hSOX2_R | GAGGAAGAGGTAACCACAGGG |
| hSFTPC_F | AGCAAAGAGGTCCTGATGGA |
| hSFTPC_R | CGATAAGAAGGCGTTTCAGG |
| h18S_F | GCAGAATCCACGCCAGTACAAG |
| h18S_R | GCTTGTTGTCCAGACCATTGGC |
| hRAGE_F | CCCTCTCCTCAAATCCACTG |
| hRAGE_R | CAGCTGTAGGTTCCCTGGTC |
| hSCGB3A2_F | GGGGCTAAGGAAGTGTGTAAATG |
| hSCGB3A2_R | CACCAAGTGTGATAGCGCCTC |
| hTP63_F | CCACAGTACACGAACCTGGG |
| hTP63_R | CCGTTCTGAATCTGCTGGTCC |
